# Phased affinity-controlled delivery of vascular endothelial growth factor, fibroblast growth factor-2, and platelet derived growth factor enhances in vitro angiogenesis

**DOI:** 10.1101/2025.07.01.662647

**Authors:** Justin E. Svendsen, Chandler L. Asnes, Samuel R. Nightheart, Madeleine R. Ford, Armaan Hajarizadeh, Simon C. Oh, Henry B. Hochstatter, Johnathan R. O’Hara-Smith, Robert E. Guldberg, Marian H. Hettiaratchi

## Abstract

Angiogenesis, the growth of vasculature from existing blood vessels, requires the coordinated secretion of multiple angiogenic growth factors that each stimulate the cellular recruitment, patterning, and morphogenesis inherent to vascular network formation. Among these secreted factors, vascular endothelial growth factor (VEGF), fibroblast growth factor-2 (FGF-2), and platelet derived growth factor (PDGF) amplify key stages of angiogenesis. Disruptions in their secretion have been implicated in poor vascular network formation. Current methods for exploring variations in the phased presentation of multiple different proteins are limited, which has restricted our ability to explore the effect of growth factor timing on angiogenesis. To address this knowledge gap, we developed affibodies, which are alpha-helical binding proteins, to phase the release of VEGF-165, FGF-2, and PDGF-BB from a single delivery vehicle via specific protein-affibody affinity interactions. We used yeast surface display to engineer three VEGF-, three FGF-2-, and two PDGF-specific affibodies with a wide range of affinities for their target proteins spanning dissociation constants of 3.08 ± 0.21 nM to 4550 ± 590 nM. We demonstrated that the cumulative release of VEGF and FGF-2 is inversely correlated with the strength of the protein-affibody affinity interaction and that hydrogels containing multiple protein-specific affibodies can control the release of VEGF, FGF-2, and PDGF, largely in accordance with the strength of the affinity interactions. Using a rat-derived intact microvascular fragment model of *in vitro* angiogenesis, we revealed that sequential delivery of soluble VEGF, followed by FGF-2, and then PDGF enhances vascular network length by 2.8-fold and branching by 4.1-fold compared to untreated MVFs. We then designed an affibody-conjugated polyethylene glycol maleimide (PEG-MAL) hydrogel to mimic this sequence of protein delivery, resulting in a 3.0-fold increase in vascular network length and a 2.3-fold increase in vascular branching compared to all other hydrogel compositions and the sequential delivery of soluble growth factors. Changing temporal growth factor presentation with affibody-conjugated hydrogels altered the expression of key angiogenic genes involved in vessel stabilization and destabilization and matrix remodeling. Perivascular coverage measured by the colocalization of lectin and alpha smooth muscle actin staining was similar between all treatment groups, suggesting pericyte recruitment to stabilize expanded vascular networks created by the soluble and affibody-mediated delivery of the optimal sequence of proteins. Overall, this work establishes a new biomaterial platform for modulating the timing of growth factor delivery, enabling the exploration of how temporal variations in protein secretion impact regeneration and development.

**Graphical Abstract:** 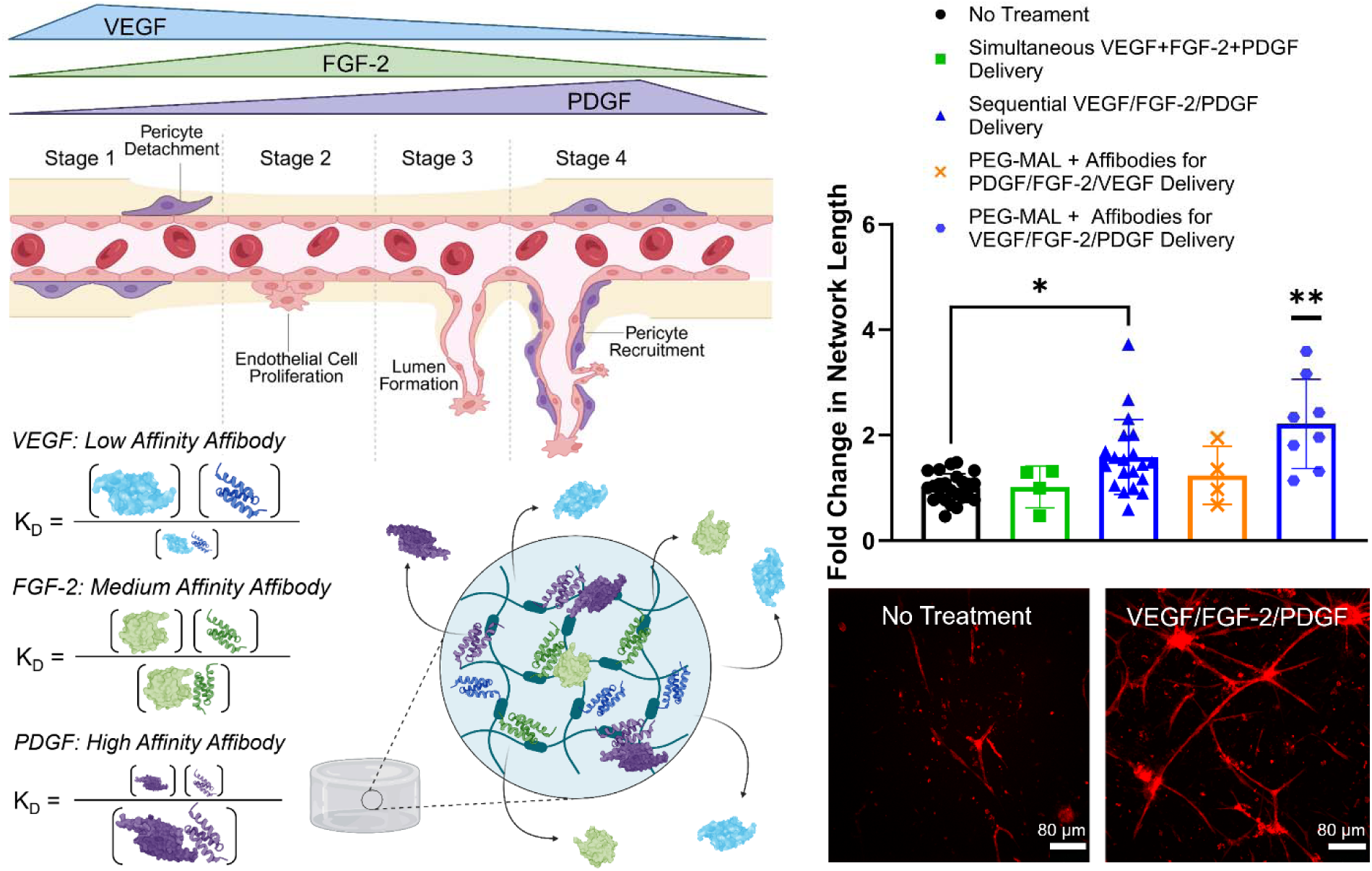

## Introduction

Coordinated sequences of biophysical and biochemical cues are required to orchestrate the cellular recruitment, patterning, and morphogenesis that occur during tissue repair.^1–4^ In the case of angiogenesis, the growth of vasculature from mature existing blood vessels, phased secretion of angiogenic growth factors such as vascular endothelial growth factor (VEGF; specifically isoform VEGF-165), fibroblast growth factor-2 (FGF-2), and platelet derived growth factor (PDGF; specifically isoform PDGF-BB) is required to coordinate several overlapping phases.^5–7^ Upon vascular injury, the angiogenic healing cascade is initiated, resulting in the secretion of a number of chemotactic, angiogenic, and other morphogenic proteins from surrounding cells.^3^ In the early stages of angiogenesis, VEGF is secreted by endothelial cells and fibroblasts proximal to the site of injury, stimulating disruption of endothelial cell-cell junctions, endothelial cell proliferation, and pericyte detachment from the lumen wall.^8^ Secreted VEGF provides both a chemotactic gradient for vascular sprouting of endothelial cells and a morphogen stimuli that differentiates stable endothelial stalk cells at the lumen wall into motile tip cells, which form a trailing neovessel.^9,10^ ^11^ Fibroblasts near the vessel secrete FGF-2, which is sequestered by the surrounding extracellular matrix.^12,13^ This local FGF-2 concentration gradient stimulates pericyte proliferation and endothelial cell chemokinesis, enabling adequate populations of pericytes for adherence to newly sprouted endothelial stalk cells and maximizing vascular coverage of endothelial cells near the site of FGF-2 enrichment.^12,14,15^ Finally, trailing neovessel stalk cells secrete PDGF, stimulating the re-adherence of local PDGF receptor beta-expressing (PDGFRB^+^) pericytes to the lumen wall to stabilize the newly formed vasculature.^16,17^ Ultimately, this coordinated sequence of cell signaling events is driven largely by the temporally regulated secretion of VEGF, FGF-2, and PDGF.^5,6^ However, chronic vascular disease and severe wounds resulting in excessive vascular damage can disrupt angiogenic protein secretion profiles, leading to dysregulated cell signaling events and impaired angiogenesis and vascular function.^14,18,19^ Consequently, methods to restore angiogenic cell signaling cascades following injury or disease may have the potential to improve vascular repair.

Exogenous protein delivery is an attractive approach for both interrogating the impact of proteins on biological systems and providing a therapeutic intervention to restore dysregulated cell signaling. Hydrogels, which are water-swollen polymer networks, have been used to deliver a variety of bioactive therapeutic proteins to different biological systems. However, hydrogels without additional modifications often rely on diffusion and material degradation to control protein release, resulting in a limited ability to sustain the presentation of physiologically relevant concentrations of therapeutic proteins at the site of interest.^20^ This approach can result in supraphysiological concentrations of therapeutic proteins rapidly diffusing from the site of interest. This rapid protein release can be especially problematic in regenerative processes such as angiogenesis where the phased release of several proteins is required to stimulate repair. Here, the burst release of multiple proteins may inhibit downstream cell signaling events, especially if at high concentrations and with proteins that have opposing biological effects.^8,21^

To address this challenge, affinity-based epitopes have been incorporated within hydrogel polymer networks to reversibly interact with target proteins and regulate their release kinetics with finer control than diffusion-mediated protein release.^22^ These affinity-based approaches include extracellular matrix molecules such as heparin that broadly interact with heparin-binding proteins, electrostatic interactions between charged proteins and materials, and DNA aptamer- or protein-based binders that have been engineered with specific affinities for target proteins.^23–26^ Engineered protein binders leverage the chemical diversity of the 20 canonical amino acids, are compatible with high-throughput cell sorting, and enable the discovery of affinity-based interactions that span sub-nanomolar to millimolar dissociation constants.^22,26,27^ Among binder scaffolds, affibodies, which are alpha-helical proteins containing 58 amino acids, have emerged as an especially promising scaffold for engineering protein-protein interactions, as they are thermally stable, mutationally tolerant, and easy to express recombinantly.^28,29^ Affibodies have previously been modified to target a variety of proteins with different affinities using both yeast surface display libraries and computational protein design.^28–30^ However, despite advances in affinity-controlled protein delivery systems, which have enabled better control over protein release kinetics, there is still a limited ability to incorporate multiple protein-specific affinity interactions into a single drug delivery vehicle to enable the phased release of multiple proteins. Thus far, the use of aptamers and affibodies in hydrogels have shown the most promise for this application.^31–33^ Since most regenerative processes involve the coordinated, temporally regulated secretion of numerous proteins, the ability to independently control the delivery of multiple proteins has the potential to both improve therapeutic approaches and enable the systematic exploration of how temporal presentation of individual proteins collectively contributes to tissue regeneration.

Here we report a library of VEGF-, FGF-2-, and PDGF-binding affibodies that can independently control the release of each target protein from polyethylene glycol maleimide (PEG-MAL) hydrogels. A yeast surface display library of affibody proteins was used to discover VEGF-, FGF-2-, and PDGF-specific affibodies with dissociation constants (K_D_) ranging from 3.07-4883 nM. We demonstrate that affibodies conjugated to PEG-MAL hydrogels control the release of target proteins with cumulative release inversely correlated to the strength of the protein-affibody affinity interaction. We establish that changing the timing of soluble VEGF, FGF-2, and PDGF delivery affects vascular network length and branching in an *in vitro* model of angiogenesis consisting of rat-derived, intact microvascular fragments. Finally, we demonstrate that mimicking the expected phased secretion of VEGF, FGF-2, and PDGF that occurs in native angiogenesis using a PEG-MAL hydrogel containing multiple protein-specific affibodies increases vascular network length and branching compared to soluble simultaneous or sequential protein delivery. Our findings indicate the potential for affibody-conjugated hydrogels to replicate the staggered protein presentation necessary to stimulate robust vascular network formation. Overall, our work establishes a foundational approach for developing highly tunable delivery systems capable of independently controlling the release of multiple angiogenic proteins and exploring the effects of their temporal regulation on angiogenesis.

## Materials & Methods

All reagents were purchased from Thermo Fisher Scientific or Sigma-Aldrich unless otherwise noted. Recombinant VEGF-165, FGF-2, and PDGF-BB were purchased from PeproTech (Rocky Hill, NJ). Biotinylated recombinant VEGF-165 and FGF-2 were purchased from Acro Biosystems (Newark, DE), and biotinylated recombinant PDGF-BB was purchased from R&D Systems (Minneapolis, MN).

### Yeast surface display

For identification of affibodies specific to VEGF-165, FGF-2, or PDGF-BB, cell sorting was performed on a yeast surface display library of the EBY100 strain of *Saccharomyces cerevisiae* containing the pCT surface display vector for galactose-inducible surface protein expression of approximately 4 × 10^8^ unique affibody sequences.^28^ Growth of the yeast surface display library and subsequent cell sorting steps were performed as previously described.^34^ Yeast were cultured in selective growth medium (16.8 g/L sodium citrate dihydrate, 3.9 g/L citric acid, 20.0 g/L dextrose, 6.7 g/L yeast nitrogen base, 5.0 g/L casamino acids) with 1 μg/mL ciprofloxacin and 100 μg/mL ampicillin in a baffled Erlenmeyer flask at 30 °C with orbital shaking at 250 rpm for 16 h. Affibody expression was induced by transferring 1 × 10^7^ cells/mL into selective induction medium (10.2 g/L sodium phosphate dibasic heptahydrate, 8.6 g/L sodium phosphate monobasic monohydrate, 19.0 g/L galactose, 1.0 g/L dextrose, 6.7 g/L yeast nitrogen base, 5.0 g/L casamino acids) with 1 μg/mL ciprofloxacin and 100 μg/mL ampicillin for an additional 16 h.

Induction of surface protein expression was verified by labeling 10^6^ yeast cells with mouse anti-c-Myc antibody (9E10, BioLegend, San Diego, CA) at 4 °C for 1 h followed by Alexa Fluor 647 goat anti-mouse IgG secondary antibody (Thermo Fisher Scientific, Waltham, MA) at 4 °C for 15 min with rotation. Cells were then washed twice with 0.1%(w/v) bovine serum albumin (BSA) in phosphate-buffered saline (PBS), and fluorescence was analyzed on an Accuri C6 Plus flow cytometer (BD Biosciences, San Jose, CA) to confirm acceptable levels of affibody surface display (30-60% of all yeast cells) prior to cell sorting.

For magnetic activated cell sorting (MACS), M-270 Carboxylic Acid Dynabeads™ (Thermo Fisher Scientific) were activated and loaded with 66 pmol of each target protein (VEGF, FGF-2, or PDGF) as previously described.^34^ Four rounds of MACS were performed for each target protein. For the first round of MACS, twenty times the clonal diversity of the induced naïve yeast library (1.72 × 10^10^ cells) were sorted. For all subsequent sorts, fifteen times the estimated clonal diversity of induced yeast from the previous positive sort was used. To perform the first negative sort, Tris-conjugated magnetic beads were combined with tubes of yeast and rotated at 4 °C for 2 h. Yeast tubes were placed against a DynaMag™-2 Magnet for 5 min to separate the yeast-bound magnetic beads from non-binding yeast. The non-binding yeast were removed, combined with BSA-conjugated magnetic beads, and rotated at 4 °C for 2 h for the second negative sort. To perform the positive target protein sort, yeast tubes were placed against the magnet for 5 min, and the supernatant yeast was removed, combined with VEGF-, FGF-2 or PDGF-conjugated magnetic beads, and stirred at 4 °C for 2 h. Following the target protein bead positive sort, yeast tubes were placed against the magnet for 5 min, the supernatant yeast was discarded, and the remaining yeast-bound beads were resuspended in growth media. Yeast from the positive sort were grown at 30 °C with orbital shaking at 250 rpm until the final concentration of the culture reached 1-10 × 10^7^ cells/mL.

For fluorescence-activated cell sorting (FACS) following MACS, 4 × 10^7^ target protein-binding yeast cells were simultaneously labeled with mouse anti-c-Myc antibody (9E10, Biolegend, San Diego, CA) and 1 μM biotinylated VEGF, FGF-2, or PDGF (bVEGF, bFGF-2, and bPDGF) by incubating at 4 °C for 1 h with rotation. Cells were washed with cold 0.1% BSA in PBS, labeled with Alexa Fluor 647 goat anti-mouse IgG secondary antibody (Thermo Fisher Scientific) and 333 nM Alexa Fluor 488 streptavidin conjugate (Thermo Fisher Scientific) by incubating at 4 °C for 15 min, and then washed twice again with 0.1% BSA in PBS. Cells were analyzed and sorted on an SH800S Cell Sorter (Sony Biotechnology, San Jose, CA). Yeast cells that were both AF-488^+^ and AF-647^+^ were sorted, expanded, and plated on agar plates for growth at 30 °C for approximately 36 h for single colony selection. Plasmid DNA from yeast was extracted, the region of interest containing the affibody sequence was amplified, and the DNA was sent to Azenta (Burlington, MA) for Sanger sequencing.

The binding affinity of each unique affibody for its target protein was characterized using flow cytometry. Yeast were stained for flow cytometry using a similar procedure to FACS staining with a few modifications. The number of induced yeast in each tube was decreased to 1 × 10^6^, solutions of 3-10,000 nM of biotinylated protein were prepared, and tubes were resuspended in 200 µL of 0.1% (w/v) BSA in PBS before being transferred to a 96-well plate. Cells were analyzed using an Accuri C6 Plus Flow Cytometer (BD Biosciences, Franklin Lakes, NJ). The equilibrium dissociation constant (K_D_) of each affibody-protein binding interaction was calculated by comparing the ratio of AF647^+^/AF488^+^ cells to AF647^+^ cells at each target protein concentration, plotting this ratio against protein concentration, and performing nonlinear regression to determine the inflection point of the curve.

### Bacterial cloning of affibody sequences

Affibody coding sequences were ordered as 300 base pair oligonucleotides from Integrated DNA Technology (IDT, Newark, NJ). Overhang primers with complementarity to the oligonucleotide 3’ and 5’ ends were designed to contain XhoI and NcoI restriction enzyme cut sites, ordered as oligonucleotides from IDT, and affibody sequences were amplified by PCR. Concurrently, pet28b+ bacterial expression vectors were plasmid extracted from previously established expression cell lines in the lab using nitrocellulose plasmid purification columns (New England Biolabs, Ipswich, MA). PCR amplified affibody-coding oligonucleotides and pET-28b+ vector were linearized by incubation with XhoI and NcoI restriction enzymes for 2 h at 37 °C. Following linearization, restriction enzymes were denatured at 60 °C for 10 min, and digested products were isolated by gel electrophoresis and subsequent gel band digestion. Bands associated with the expected digested product molecular weights were excised, and DNA was harvested by standard DNA-gel extraction (New England Biolabs). Gel extracted products were co-incubated with T4 ligase in T4 ligation buffer for 1 h to conjugate complimentary overhangs from restriction enzyme digestion. T4 ligated products were drop dialyzed on 0.22 µm nitrocellulose filters for 30 min, and ligated affibody coding pET-28b+ vectors were cloned into chemically competent DH5alpha *E. coli* for plasmid storage and BL21 (DE3, New England Biolabs) *E. coli* for protein expression. Transformed cells were screened under antibiotic selection on LB agar plates containing 50 µg/mL kanamycin. Colonies were isolated, plasmid extracted and sent for full backbone sequencing through Plasmidsaurus (Eugene, OR) to verify successful incorporation of affibody coding pET-28b+ vector.

### Bacterial expression and purification of affibodies

To verify protein expression from cloned *E. coli*, small scale protein expression was performed as previously described.^35^ For large scale protein expression, 10 mL of LB with 50 µg/mL kanamycin were inoculated with swabs from bacterial glycerol stocks and incubated for approximately 16 h at 37°C and 220 rpm in a shaking incubator. In parallel, terrific broth (TB) (Research Products International, Mount Prospect, IL) was prepared by adding 85.7 g of TB powder and 14.4 mL of 50% (w/v) glycerol to 1.8 L of double distilled water and autoclaved for sterilization. On the next day, 1.8 L TB were supplemented with 50 µg/mL of kanamycin and 10-15 drops of antifoam agent (Antifoam 204) and inoculated with the 10 mL bacteria cultures from the previous day. TB cultures were incubated at 37°C in a LEX-10 water bath bioreactor (Epiphyte, Toronto, ON, Canada) until an optical density at 600 nm (OD_600nm_) greater than 0.8 was achieved. Next, protein expression was induced with the addition of 0.5 µM isopropyl β-D-1-thiogalactopyranoside (IPTG) (GoldBio, St. Louis, MO) for 14-18 h at 18°C. Bacterial cultures were then pelleted by ultracentrifugation at 6000 rpm at 4°C for 20 min and stored at -80°C.

Bacteria pellets were thawed into 35 mL binding buffer (5 mM imidazole, 500 mM NaCl, 50 mM Tris (GoldBio) containing 8.6 mM tris(2-carboxyethyl)phosphine HCl (TCEP; GoldBio) and sonicated on ice at 55% amplitude for 5 cycles (15 s on, 45 s off). Bacterial lysate was subsequently centrifuged at 13,000 rpm and 4°C for 30 min, and supernatant was collected for immobilized metal affinity chromatography. Supernatant was added to 1.8 mL of cobalt agarose beads (GoldBio) and rotated at 4°C for approximately 4 h. The agarose supernatant mixtures were added to glass chromatography columns and washed 5 times with 10 mL of wash buffer containing TCEP (50 mM Tris, 500 mM NaCl, 30 mM imidazole, 10 mM TCEP), followed by 5 washes with 10 mL of wash buffer without TCEP. Affibodies were then eluted with approximately 15 mL of elution buffer (50 mM Tris, 500 mM NaCl, 250 mM imidazole) in 1 mL increments until all protein was eluted, as confirmed by Bradford reagent. Eluted protein was buffer exchanged into PBS and concentrated to 1-5 mg/mL using a 3 kDa molecular weight cut off (MWCO) centrifugal filter (Millipore, Burlington, MA) and stored at -80°C.

Size exclusion chromatography was performed using a HiPrep 16/60 200 HR chromatography column (Cytiva, Marlborough, MA) on an NGC Chromatography System (Bio-Rad Laboratories, Hercules, CA). The column was equilibrated with degassed PBS. Affibody samples were thawed and filtered through 0.22 µm filters to remove any insoluble particulates from solution. The sample was loaded onto the column, and 2 mL fractions were collected along a 180 mL elution range with affibody-containing fractions collected based on 280 nm absorbance signal. Sodium dodecyl sulfate– polyacrylamide gel electrophoresis (SDS-PAGE) followed by Coomassie Brilliant Blue staining (Bio-Rad Laboratories) was used to visualize protein bands and confirm affibody purity in the fractions of interest. Fractions were then pooled, concentrated using 3 kDa MWCO centrifugal filters, aliquoted, and stored at -80 C.

### Biolayer interferometry

Binding specificity of each soluble VEGF-, FGF-2-, and PDGF-specific affibody to their respective target protein was confirmed by biolayer interferometry (BLI) using a GatorPlus biolayer interferometer (GatorBio, Palo Alto, CA). Biotinylated VEGF, FGF-2, or PDGF was diluted to 25 nM in 0.05% (v/v) Tween 20 in PBS (PBST) and loaded onto streptavidin-coated glass BLI probes for 180 s until a wavelength shift of approximately 1 nm was achieved. Probes were subsequently submerged in PBST 300 seconds to remove any loosely bound protein. Probes were then incubated in 1000 nM of either VEGF-, FGF-2-, or PDGF-specific affibodies for 600 s to measure protein-affibody association and subsequently moved to PBST for 600 s to measure protein-affibody dissociation. Three control probes were loaded with bVEGF, bFGF-2, or bPDGF only, and additional control probes without growth factors underwent association with 1000 nM of each affibody. These control probes were used to subtract out signal drift and non-specific binding of affibodies to the probes, respectively.

### Circular dichroism

Secondary structures of affibodies were confirmed by circular dichroism using a Jasco J-815 spectropolarimeter (Jasco, Easton, MD) as previously described.^35^ Isoelectric points of affibodies were calculated using Expasy to select buffer conditions with a minimum pH difference of 0.5 from the affibody isoelectric point to avoid protein unfolding. Purified affibodies were buffer exchanged into 10 mM Tris pH 7.4 using 3 kDa MWCO centrifuge filters and diluted to 0.3 mg/mL. All affibody samples were loaded into 2 mL 0.1 cm quartz cuvettes (Starna, Atascadero, CA) for data acquisition. High tension voltage and absorption spectra were measured across 190-250 nm in triplicate at 10 nm/min using a step size of 1 nm at room temperature. Triplicate absorption spectra were normalized, and background signal from the buffer was subtracted prior to data analysis. Signal measurements (mdeg) were converted into molar ellipticity and normalized to affibody molecular weight and concentration.

### Affibody-conjugated hydrogel synthesis and protein release

100 µL single affibody-conjugated polyethylene glycol maleimide (PEG-MAL) hydrogels were synthesized in 2.0 mL low retention microcentrifuge tubes as previously described.^34,35^ VEGF-, FGF-2-, or PDGF-specific affibodies were added to 20 kDa 4-arm PEG-MAL (Laysan Bio, Arab, AL) suspended in PBS pH 7.4 at a 500:1 molar ratio of affibodies to growth factor and incubated for 2 h at 4 °C. The C-terminal cysteine on the affibodies reacted with the maleimides on the PEG-MAL through a Michael-type addition reaction. 5% (w/v) affibody-conjugated PEG-MAL was added to centrifuge tubes with ∼ 74 µg dithiothreitol (DTT) in PBS to occupy all remaining maleimide groups and crosslinked for 1 h at 4 °C. Following crosslinking, hydrogels were swelled overnight with 1.8 mL PBS pH 7.4 at 4 °C with gentle rotation. Swelled hydrogels were washed with 4 mL of fresh PBS and loaded overnight with 100 ng of VEGF, FGF-2, or PDGF in a low volume (20 µL) of 0.1% (w/v) BSA in PBS at 4 °C with orbital shaking. Supernatant was then collected to calculate the amount of protein encapsulated prior to protein release. Protein release was initiated upon addition of 900 µL of 0.1% (w/v) BSA in PBS, and the hydrogels were incubated at 37 °C for 7 days. Timepoints were taken at 0, 15 min, 30 min, 1 h, 3 h, 6 h, and 1, 2, 3, 4, 5, 6, and 7 days by removal of 200 µL of supernatant, and replacement with 200 µL of fresh 0.1% (w/v) BSA in PBS. Protein concentrations were measured using protein-specific enzyme-linked immunosorbent assays (ELISA, PeproTech). Cumulative protein release was plotted over 7 days as a percentage of initial encapsulated protein.

Multiple affibody-conjugated PEG-MAL hydrogels were synthesized similarly to contain combinations of VEGF-, FGF-2-, and/or PDGF-specific affibodies with varying affinities. Hydrogels were synthesized as described above with 500:1 molar ratios of affibody to loaded growth factors. Following overnight swelling in PBS pH 7.4, hydrogels were loaded with 8.75 pmols each of VEGF, FGF-2, and PDGF in a low volume (20 µL) of 0.1% (w/v) BSA in PBS. Protein was released over 7 days into complete microvascular fragment minimal media. Protein encapsulation and release were quantified by ELISA.

### Endothelial tube formation assay

Human umbilical vein endothelial cells (HUVECs, ATCC, Manassas, VA) were seeded in tissue culture flasks at 2,500 cells/cm^2^ and cultured in complete endothelial growth medium (Lonza, Walkersville, MD) at 37° C and 5% CO_2._ Upon reaching 60-80% confluence, cells were washed with PBS, trypsinized, and seeded at 5,000 or 15,000 cells/well onto 96-well plates pre-coated with 100 µL of Cultrex^TM^ reduced growth factor basement membrane extract (R&D Systems). HUVECs were allowed to adhere to the coated surface for 2 h. 10 ng/mL of VEGF, FGF-2, and/or PDGF were then added either separately or simultaneously to the wells to assess the effects of the growth factors on HUVEC network length and branching. Cells were cultured for 48 h on a BioTek Lionheart FX automated microscope (Agilent Technologies, Santa Clara, CA) at 37° C and 5% CO_2_ with 4x phase contrast images taken at 4, 8, 12, 16, 20, 24, and 48 hours. Total network length and branching of HUVEC networks were measured from images using the Angiogenesis Analyzer plugin for ImageJ.^36^

### Microvascular fragment (MVF) isolation, culture, and media collection

All surgical procedures were conducted according to the University of Oregon Institutional Animal Care and Use Committee protocol for MVF harvest. MVFs were isolated from epididymal fat pads of retired breeder Lewis rats (> 350 grams) as previously described, with minor modifications.^37^ Harvested tissues were manually minced and further digested via hand mixing in a solution of 2.3 mg/mL collagenase type 1 (Worthington, Lakewood, NJ), 1.3 mg/mL DNase I, and 5 mg/mL BSA in a 37 °C water bath. The digested tissue was then centrifuged to separate out undigested matrix and washed three times with Hank’s buffered saline solution (HBSS) supplemented with 5% (v/v) heat-inactivated fetal bovine serum (FBS). The digested tissue was then resuspended in FBS-HBSS and filtered sequentially through 200 and 20 µm nylon meshes to remove undigested matrix, larger fragments, and single cells to isolate fragments between 20-200 µm in size. After filtering, fragments were counted via brightfield imaging in 20 µL volumes to approximate total yield and assessed for viability using a NucleoCounter NC-200 (Chemometec, La Jolla, CA). Filtered MVFs were resuspended at 20,000 fragments/mL in a solution of 0.3% (w/v) rat collagen type I in Dulbecco’s Modified Eagle Medium (DMEM) supplemented with 62 mM N-(2-hydroxyethy)piperazine-N’-(2-ethanesulfonic acid) (HEPES) and 226 mM sodium bicarbonate. The collagen solution containing MVFs was then pipetted into either a 24- or 96-well plate, allowed to form a hydrogel, and cultured in serum-free DMEM/F-12 medium containing 100 µg/mL apo-transferrin, 100 µg/mL BSA, 10 µg/mL insulin, 100 µM putrescine, 30 nM sodium selenite, 20 nM progesterone, and 1% (v/v) penicillin-streptomycin. A stable MVF culture with minimal cell death was achieved after 3 days, as previously established.^38,39^ The MVFs were cultured at 37 °C and 5% CO_2_ for 7 days, beyond which point the collagen hydrogel was susceptible to degradation by proteases secreted by the MVFs under these treatment conditions.^40^ Media were changed on days 3 and 5. Soluble growth factor treatments were delivered on days 3, 4, and 5. For treatment with affibody-conjugated hydrogels, 200 µL 5% (w/v) PEG-MAL hydrogels containing combinations of VEGF-, FGF-2-, and/or PDGF-specific affibodies and 17.5 pmols of each growth factor were synthesized in 24-well transwell inserts with 1 μm pores, washed to remove excess DTT, and transferred to MVF plates on day three of culture. By fabricating the PEG-MAL hydrogels in transwell inserts, we physically separated the protein delivery hydrogel from the MVFs in the collagen hydrogel. After 7 days, conditioned media samples were collected to measure VEGF, FGF-2, and PDGF concentrations using protein-specific ELISAs. The collagen hydrogels were then fixed with 4% (v/v) paraformaldehyde and stained with 20 µg/mL of rhodamine-labeled Griffonia (Bandeiraea) Simplicifolia Lectin I (Vector Laboratories, Newark, CA). Z- stack images (250 µm depth with 5 µm step size) were captured using a CSU-W1 SoRa Spinning Disk confocal microscope (Nikon, Melville, NY). MVF experiments were performed with tissue isolated from a total of 42 rats over 8 independent harvests with at least 4 biological replicates per treatment group.

### Volumetric image analysis

Maximum projection z-stack confocal images of MVFs were processed as previously described using Amira Software (Thermo Fisher Scientific) with deconvolution, median filtering, and a z-drop correction.^41^ Briefly, clusters smaller than 60 voxels were removed using Amira’s “Remove Islands” module, and remaining volumes were segmented and skeletonized. Network length and branching were analyzed using a centerline tree algorithm. Short fragments (<200 µm) which failed to grow were excluded from the analysis. Vascular network length and branching for each treatment group were normalized to the mean network length and branching of untreated MVFs grown in standard culture medium.

### Perivascular recruitment

After 7 days of culture, lectin-stained MVFs were also stained for the presence of alpha-smooth muscle actin (α-SMA) to determine the degree of perivascular coverage. 400 μL collagen hydrogels were permeabilized with 400 μL of 0.1% Triton X-100 and 1 mM CaCl_2_ in PBS for 15 minutes at room temperature, followed by three PBS washes for 10 min each. Hydrogels were then blocked with 400 μL of 5% (w/v) BSA in PBS for 2 h at room temperature. All staining steps were performed in the dark. Staining with the primary anti-mouse α-SMA antibody (1:200 in 1% (w/v) BSA in PBS, ab7817, Abcam, Cambridge, UK) was performed overnight at 4 °C, followed by three PBS washes for 10 minutes each. Staining with the secondary goat-anti-mouse Alexa Fluor 647 antibody (1:500 in PBS) was performed for 30 min at 4 °C, followed by three PBS washes for 10 minutes each. Hydrogels were then stored in PBS at 4 °C until imaging.

Z-stack images (250 µm depth with 5 µm step size) were captured using a CSU-W1 SoRa Spinning Disk confocal microscope. Confocal z-stacks were processed using ImageJ with a custom macro. Five slices of the 50-slice stack were isolated for analysis: slice 5, 15, 25, 35, and 45. Each slice was cropped to 2048×1968 pixels to exclude edge artifacts. Local contrast enhancement was applied using contrast limited adaptive histogram equalization (CLAHE), and noise was reduced via median filtering. α-SMA (blue) and lectin (red) signals were separately auto-thresholded using Otsu’s algorithm to generate binary masks. Both channels were then analyzed using the Colocalization Threshold plugin to calculate Mander’s overlap coefficients (M1), representing the percentage of α-SMA signal colocalized with lectin-positive area. Within each representative stack, the individual M1 coefficients for each slice were averaged and reported as a representative value for the stack.

### Gene expression analysis

Real time quantitative polymerase chain reaction (RT-qPCR) was performed to assess the gene expression of MVFs treated with single soluble growth factors, multiple soluble growth factors, or multiple growth factors released from different PEG-MAL hydrogel compositions. Collagen hydrogels containing MVFs were digested on Day 7 by the addition of 500 μL of TRIzol^TM^ (Fisher Scientific, Waltham, MA), followed by the addition of 200 μL of 100% (v/v) chloroform to trigger phase separation. Phase-separated fractions were centrifuged at 15,000 rpm for 5 minutes, and the aqueous top layers were collected to isolate RNA from lipids, proteins, and other cell debris. Collected aqueous RNA fractions received 700 μL of 70% (v/v) ethanol and were briefly agitated to ensure complete mixing. RNA suspensions were then collected using spin columns according to manufacturer’s instructions (PureLink RNA extraction kit; Invitrogen, Carlsbad, California). Extracted RNA concentrations were determined by UV-vis spectroscopy (Implen 80), and RNA quality was confirmed by A260/A280 > 1.6 and A260/A230 > 1.8 readings. cDNA synthesis was performed according to manufacturer’s instructions (LunaScript cDNA reaction kit; New England Biolabs, Ipswich, MA) with 250 ng of RNA per reaction. cDNA was stored at -80 °C until use.

All RT-qPCR was performed using 1:10 dilutions with three technical replicates for each biological replicate. TaqMan_™_ probes for the following target genes were ordered from Thermo Fisher Scientific: matrix metalloproteinase 3 (MMP-3, Rn00591740_m1), tissue inhibitor of metalloproteinases 1 (TIMP-1, Rn00587558_m1), angiopoietin 1 (ANGPT-1, Rn01504818_m1), angiopoietin 2 (ANGPT-2, Rn01756774_m1), VEGF A (VEGF-A, Rn01511602_m1), VEGF receptor 2 (VEGFR-2, Rn00564986_m1), PDGF receptor beta (PDGFRΒ, Rn00709573_m1), and beta actin (ACTB, Rn00667869_m1) as a housekeeping gene. Housekeeping gene stability was verified across all treatment conditions with < 0.5 difference in threshold cycles (Ct) between all treatment conditions. The standard deviation between technical replicates of threshold cycles was confirmed to be < 0.5 Ct. All data were collected and analyzed using QuantStudio_™_ 5 Real Time PCR System and associated QuantStudio_™_ Design and Analysis Software package (Thermo Fisher Scientific). Changes in relative gene expression were calculated using the 2^-ΔΔCt^ method and plotted as fold change in gene expression compared to untreated MVFs on Day 7.

### Statistical analysis

Unless otherwise noted, all data were analyzed and graphed using GraphPad Prism version 10.0.3 (Boston, MA). Ordinary one- or two-way ANOVAs with Tukey’s multiple comparison test were used to determine statistical differences. Dissociation constants of yeast-displayed affibodies binding to target proteins were calculated using non-linear regression curve fitting with a one site-specific binding model and reported as mean ± standard error of the mean. Unless otherwise noted, all other data are presented as mean ± standard deviation. p < 0.05 was considered statistically significant.

## Results

### Yeast-surface display enables discovery of VEGF-, FGF-2-, and PDGF-specific affibodies

To identify affibodies with a broad range of affinities towards VEGF, FGF-2, and PDGF, we first screened a yeast surface display library containing 400 million affibody variants using MACS and FACS. We identified two VEGF-specific affibodies with dissociation constants of 58.3 ± 13.7 nM and 307 ± 52.68 nM, which were named VEGF High Affibody and VEGF Medium Affibody, respectively (**Fig. 1A**, **1B**). To obtain a lower affinity VEGF-specific affibody to achieve the desired fast VEGF release, we used site-directed mutagenesis to mutate the aspartic acid at residue 32 to an alanine on a previously reported VEGF-specific affibody,^35^ resulting in a VEGF Low Affibody with a dissociation constant of 4186 ± 1706 nM, which is an order of magnitude lower than that of the VEGF Medium Affibody. We identified three FGF-2-specific affibodies with dissociation constants of 3.08 ± 0.21 nM, 121.2 ± 16.8 nM, and 4550 ± 590 nM, similarly spanning three orders of magnitude and named FGF-2 High Affibody, FGF-2 Medium Affibody, and FGF-2 Low Affibody, respectively (**Fig. 1C**, **1D**). We identified one previously published PDGF-specific affibody with a dissociation constant of 855 ± 255 nM using yeast surface display,^35^ which we define here as a PDGF Medium Affibody. To obtain additional PDGF-specific affibodies with different affinities and high specificity for PDGF, we used the computational protein design tool Rosetta to mutate our PDGF Medium Affibody, resulting in three PDGF specific affibodies (PDGF Affibody-11, PDGF-Affibody-13, and PDGF-Affibody-16) with dissociation constants of 6.44 nM, 77.4 nM, and 5.87 nM, respectively.^35^ We moved forward with the computationally designed PDGF Affibody-13, which we define here as a PDGF High Affibody.

**Figure 1.**
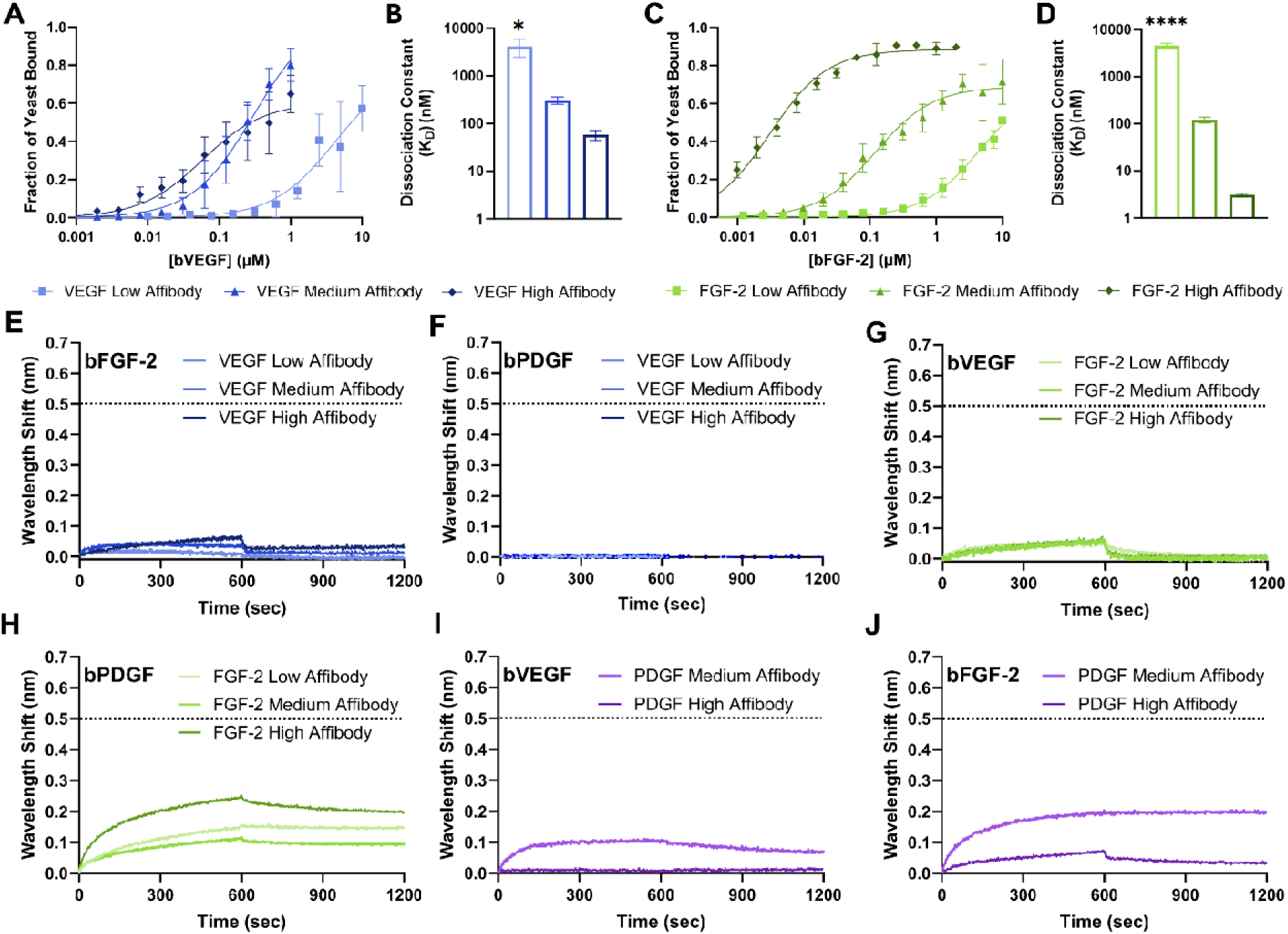
Flow cytometry and biolayer interferometry analysis of affibodies binding to on-target and off-target proteins. A) Fraction of yeast displaying VEGF-specific affibodies binding to 3-10,000 nM bVEGF. (n=3) The following dissociation constants were determined for B) VEGF binding to VEGF-specific affibodies: VEGF High Affibody K_D_ = 58.3 ± 13.8 nM, VEGF Medium Affibody K_D_ = 306.9 ± 52.7 nM, VEGF Low Affibody K_D_ = 4186 ± 1706 nM. C) Fraction of yeast displaying FGF-2-specific affibodies binding to 3-10,000 nM bFGF-2. (n=3) The following dissociation constants were determined for D) FGF-2 binding to FGF-2-specific affibodies: FGF-2 High Affibody K_D_ = 3.08 ± 0.21 nM, FGF-2 Medium Affibody K_D_ = 121 ± 16.8 nM, FGF-2 Low Affibody K_D_ = 4550 ± 590 nM. For BLI, 25 nM bVEGF, bFGF-2, or bPDGF were loaded onto streptavidin-functionalized probes and allowed to associate 1000 nM of either VEGF-, FGF-2-, or PDGF-specific affibodies. BLI sensograms depicting binding between E) VEGF-specific affibodies to bFGF-2, F) VEGF-specific affibodies to bPDGF, G) FGF-2-specific affibodies to bVEGF, H) FGF-2-specific affibodies to bPDGF, I) PDGF-specific affibodies to bVEGF, and J) PDGF-specific affibodies to bFGF-2. Dotted line indicates typical saturating wavelength shift of 0.5 nm for specific affinity interactions. Statistical significance was determined by one-way ANOVA and Tukey’s post-hoc test. (n=4, *p < 0.05, ****p<0.0001)

### Biochemical characterization of VEGF-, FGF-2-, and PDGF-specific affibodies

VEGF-, FGF-2-, and PDGF-specific affibodies were cloned into pet28b+ overexpression vectors with a c-terminal adjacent hexa-histidine tag for protein purification using immobilized metal affinity chromatography and a c-terminal cysteine for conjugation to hydrogels. Recombinant affibodies were purified from sonicated bacterial lysate using immobilized metal affinity chromatography followed by size exclusion chromatography. All affibodies displayed the expected molecular weights of ∼ 7kDa on SDS-PAGE **(Fig. S1A)** and high degrees of alpha-helical folding as demonstrated by far-UV circular dichroism molar ellipticity **(Fig. S1B-D)**.

To determine the specificity of VEGF-, FGF-2, and PDGF-specific affibodies for their target proteins, we used biolayer interferometry (BLI) to measure off-target binding between recombinantly expressed soluble affibodies and their target proteins. Our previous work reporting VEGF and PDGF binding to their respective affibodies demonstrated a repeatable saturating wavelength shift of 0.5 nm for specific affinity interactions,^35^ providing a point of comparison for interpreting the degree of non-specific binding observed herein under the same experimental conditions. All VEGF-specific affibodies displayed negligible binding to both bFGF-2 and bPDGF **(Fig. 1E**, **1F)**. All FGF-2 affibodies displayed negligible binding to bVEGF and moderate binding to bPDGF, with the FGF-2 High Affibody displaying the highest off-target binding to bPDGF **(Fig. 1G**, **1H)**. Our previous work computationally modelling the binding interface between the PDGF High Affibody and PDGF-BB predicted affibody binding at the PDGF receptor beta (PDGFRB) epitope on PDGF-BB.^35^ This epitope has been implicated in mediating the heparin binding capacity of PDGF-BB and may be complimentary to the heparin binding domain of FGF-2, permitting the observed off-target binding between PDGF and the FGF-2 High Affibody.^42^ Finally, the PDGF High Affibody displayed no binding to bVEGF or bFGF-2, while the PDGF Medium Affibody exhibited moderate binding to bVEGF and bFGF-2 (**Fig. 1I**, **1J**). This off-target binding may be described by the shared capacity of FGF-2 and PDGF-BB to bind heparin with conserved heparin binding domains.^43–45^

### VEGF- and FGF-2-specific affibodies control protein release from hydrogels according to affibody affinity

We have previously demonstrated that our PDGF-specific affibodies can control the delivery of PDGF from PEG-MAL hydrogels, with the amount of PDGF released being inversely correlated with the affinity of the affibody for PDGF.^48^ We next explored the impact of VEGF- and FGF-2-specific affibodies on controlling growth factor delivery from PEG-MAL hydrogels. PEG-MAL hydrogels were conjugated with VEGF- or FGF-2-specific affibodies using thiol-ene click chemistry between the affibody c-terminal cysteine and the PEG-MAL maleimide alkene at a molar ratio of 500:1 affibodies to protein. VEGF and FGF-2 were soaked into the hydrogels overnight, and the unencapsulated protein was removed to quantify initial protein encapsulation. All hydrogels displayed similar VEGF encapsulation (>65%), regardless of the presence of VEGF affibodies **(Fig. 2A)**. Hydrogels conjugated with the VEGF High Affibody released less VEGF over 7 days (52.0 ± 8.4%) than all other compositions, with significant differences starting at 6 hours **(Fig. 2B)**. In contrast, the hydrogel without affibodies released 74.5 ± 9.0% of the loaded VEGF, the VEGF Low Affibody hydrogel released 74.6 ± 8.6% of the loaded VEGF, and the VEGF Medium Affibody hydrogel released 88.8 ± 7.5% of the loaded VEGF. These protein release outcomes were as expected as the VEGF High Affibody displayed approximately 5-fold and 100-fold greater affinities for VEGF than the VEGF Medium and VEGF Low Affibodies, respectively. However, no significant differences were observed at any timepoint between hydrogels without affibodies, hydrogels with the VEGF Medium Affibody, and hydrogels with the VEGF Low Affibody, indicating that these affinity interactions were not strong enough to reduce VEGF release at these VEGF concentrations and affibody molar excess. Thus, we chose to move forward with only the VEGF High Affibody for multiple growth factor delivery.

**Figure 2.**
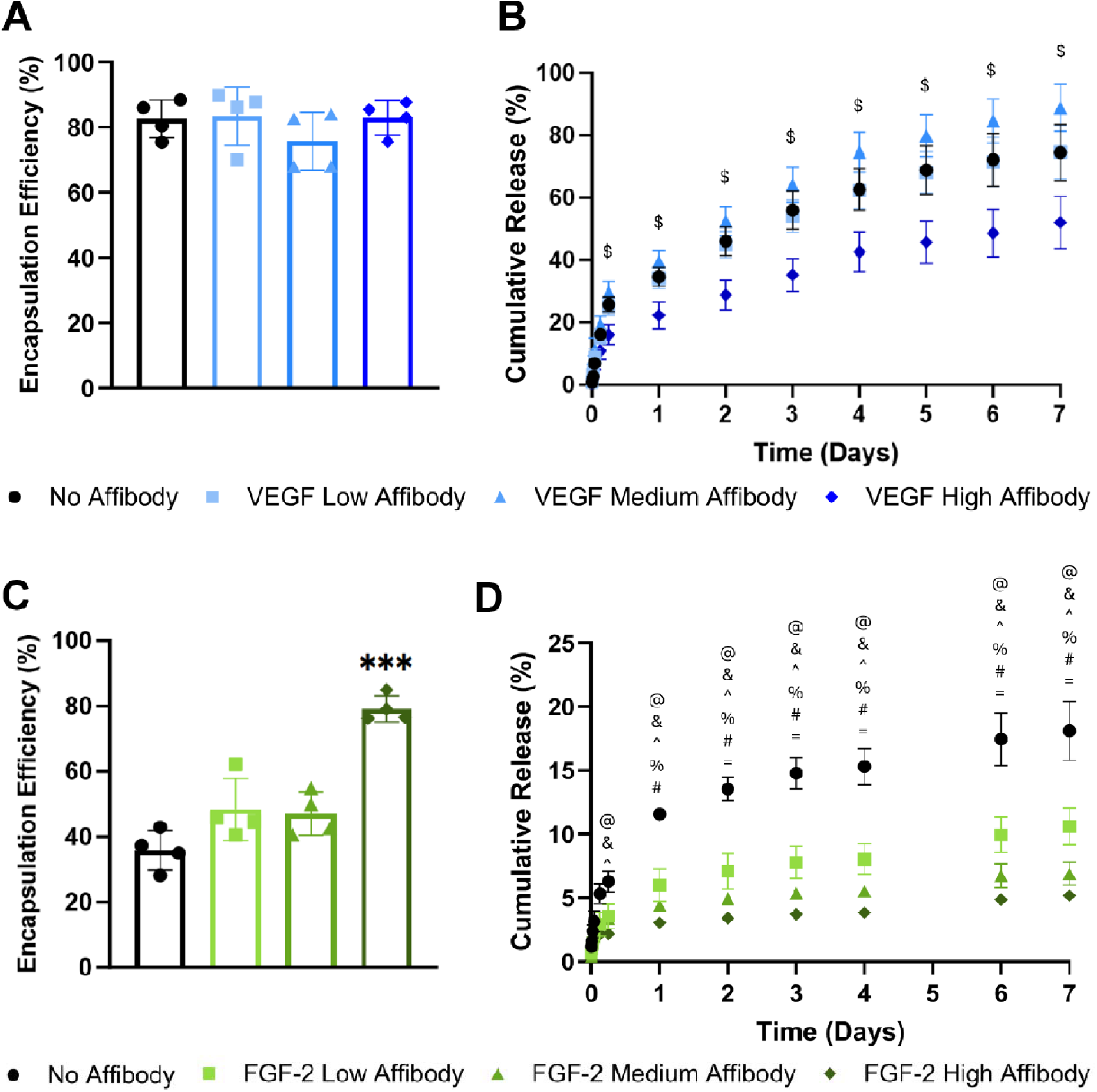
Controlled release of VEGF and FGF-2 from single affibody-conjugated hydrogels. A) Encapsulation efficiency of VEGF loaded within VEGF-specific affibody-conjugated hydrogels. Statistical significance determined by one-way ANOVA and Tukey’s post hoc test. (n=4) B) Controlled release of VEGF into 0.1% (w/v) BSA in PBS at 37 °C over 7 days from VEGF-specific affibody-conjugated hydrogels. Statistical significance determined by two-way ANOVA and Tukey’s post hoc test. (n=4, $ p < 0.05 for VEGF High Affibody compared to all other groups) C) Encapsulation efficiency of FGF-2 loaded within FGF-2-specific affibody-conjugated hydrogels. Statistical significance determined by one-way ANOVA and Tukey’s post hoc test. (n=4, ***p<0.001) D) Controlled release of FGF-2 into 0.1% (w/v) BSA in PBS at 37 °C over 7 days from FGF-2-specific affibody-conjugated hydrogels. Statistical significance determined by two-way ANOVA and Tukey’s post hoc test. (n=4, *p < 0.05, @ No Affibody vs. FGF-2 Low Affibody, & No Affibody vs. FGF-2 Medium Affibody, ^ No Affibody vs. FGF-2 High Affibody, % FGF-2 Low Affibody vs. FGF-2 Medium Affibody, # FGF-2 Low Affibody vs. FGF-2 High Affibody, = FGF-2 Medium Affibody vs. FGF-2 High Affibody)

PEG-MAL hydrogels containing the FGF-2 High Affibody displayed higher FGF-2 encapsulation than all other hydrogel compositions, consistent with the expected impact of incorporating a nanomolar affinity binder into the polymer network **(Fig. 2C)**. All affibody-containing hydrogels displayed lower release of FGF-2 than the hydrogels without affibodies. Furthermore, the amount of FGF-2 released over 7 days was inversely correlated with the affinity strength of the incorporated FGF-2 affibody, with the hydrogel without affibodies releasing 18.1 ± 2.3%, the FGF-2 Low Affibody hydrogel releasing 10.6 ± 1.4%, the FGF-2 Medium Affibody hydrogel releasing 6.9 ± 0.9%, and the FGF-2 High Affibody hydrogel releasing 5.2 ± 0.4% **(Fig. 2D)**. These results are consistent with FGF-2-specific affibody affinities for FGF-2, with the FGF-2 High Affibody displaying approximately 40-fold and 1500-fold greater affinities for FGF-2 than the FGF-2 Medium Affibody and FGF-2 Low Affibody, respectively. Overall, less FGF-2 was detected over 7 days than VEGF, which is likely due to the short half-life of FGF-2 in solution at 37°C.^49,50^ In both cases, not all of the loaded growth factor was retrieved from the hydrogels. This is consistent with other studies involving affinity-controlled protein release,^32,51,52^ in which the remaining protein is likely still bound to the affibodies in the hydrogel or undergoes denaturation that makes it undetectable via ELISA.

### Multiple protein-specific affibodies control VEGF, FGF-2, and PDGF release from hydrogels

With the goal of co-delivering VEGF, FGF-2, and PDGF from a single delivery vehicle, we next sought to characterize protein release from PEG-MAL hydrogels containing multiple protein-specific affibodies. Instead of evaluating every possible combination of low-, medium-, and high-affinity affibodies for each target protein, we focused on combinations of affibodies that would allow us to best investigate the temporal roles of VEGF, FGF-2, and PDGF on angiogenesis. This included a PEG-MAL + Affibodies (Optimal) composition inspired by the relative roles of VEGF, FGF-2, and PDGF at each stage of angiogenesis, in which VEGF initiates the initial vascular destabilization stage of angiogenesis,^8,11^ followed by FGF-2-induced vascular remodeling,^12,14^ and then finally PDGF induced re-stabilization^5,6,8,10^; a PEG-MAL + Affibodies (High-Affinity) composition, in which high-affinity affibodies are incorporated into the hydrogel to release the minimal amount of each target protein; and finally, a PEG-MAL + Affibodies (Reverse) composition in which the proteins are released with the reverse timing of what is typically observed during angiogenesis, with PDGF released first, followed by FGF-2, and then VEGF last. We also compared these hydrogel compositions to a PEG-MAL Simultaneous hydrogel without affibodies. Based on our yeast surface display results for affinity strengths, BLI results for affibody specificity, and endpoint cumulative protein release, we chose to move forward with the VEGF High Affibody, FGF-2 Medium Affibody, FGF-2 High Affibody, and PDGF High Affibody to achieve the desired release profiles for each growth factor. We used different combinations of affibodies in each hydrogel formulation to achieve target release profiles **(Table 1)**. PEG-MAL hydrogels were conjugated with combinations of VEGF-, FGF-2-, and PDGF-specific affibodies at 500:1 molar ratios of affibodies to growth factors. Hydrogels were loaded with 17.5 pmols of each growth factor to achieve protein concentrations that would stimulate cellular responses within the initial 3 days of protein release.^2,3^

**Table 1.** Compositions of hydrogels containing multiple protein-specific affibodies.

|  | **VEGF** | **FGF-2** | **PDGF** |
| --- | --- | --- | --- |
| **PEG-MAL Simultaneous** | No VEGF affibody | No FGF-2 affibody | No PDGF affibody |
| **PEG-MAL + Affibodies (Optimal)** | No VEGF affibody | FGF-2 Medium Affibody | PDGF High Affibody |
| **PEG-MAL + Affibodies (High-Affinity)** | VEGF High Affibody | FGF-2 High Affibody | PDGF High Affibody |
| **PEG-MAL + Affibodies (Reverse)** | VEGF High Affibody | FGF-2 Medium Affibody | No PDGF Affibody |

After loading with growth factors, unencapsulated protein was removed from solution and quantified, and the growth factor-loaded hydrogels were allowed to release protein into MVF minimal cell medium at 37 °C for 7 days. Unexpectedly, the PEG-MAL + Affibodies (Reverse) hydrogels encapsulated less VEGF than all other groups, despite containing the same VEGF High Affibody as the PEG-MAL + Affibodies (High-Affinity) hydrogels, suggesting interactions between the multiple types of affibodies and growth factors in the hydrogel **(Fig. 3A)**. The PEG-MAL + Affibodies (High-Affinity) hydrogels containing the FGF-2 High Affibody encapsulated more FGF-2 than all other groups **(Fig. 3B)**, which was consistent with the higher FGF-2 encapsulation observed for single protein hydrogels containing the FGF-2 High Affibody **(Fig. 3C)**. As expected, the PEG-MAL + Affibodies (High-Affinity) hydrogels containing the PDGF High Affibody also encapsulated more PDGF than the PEG-MAL + Affibodies (Reverse) hydrogels that contained no PDGF-specific affibodies **(Fig. 3C)**.

**Figure 3.**
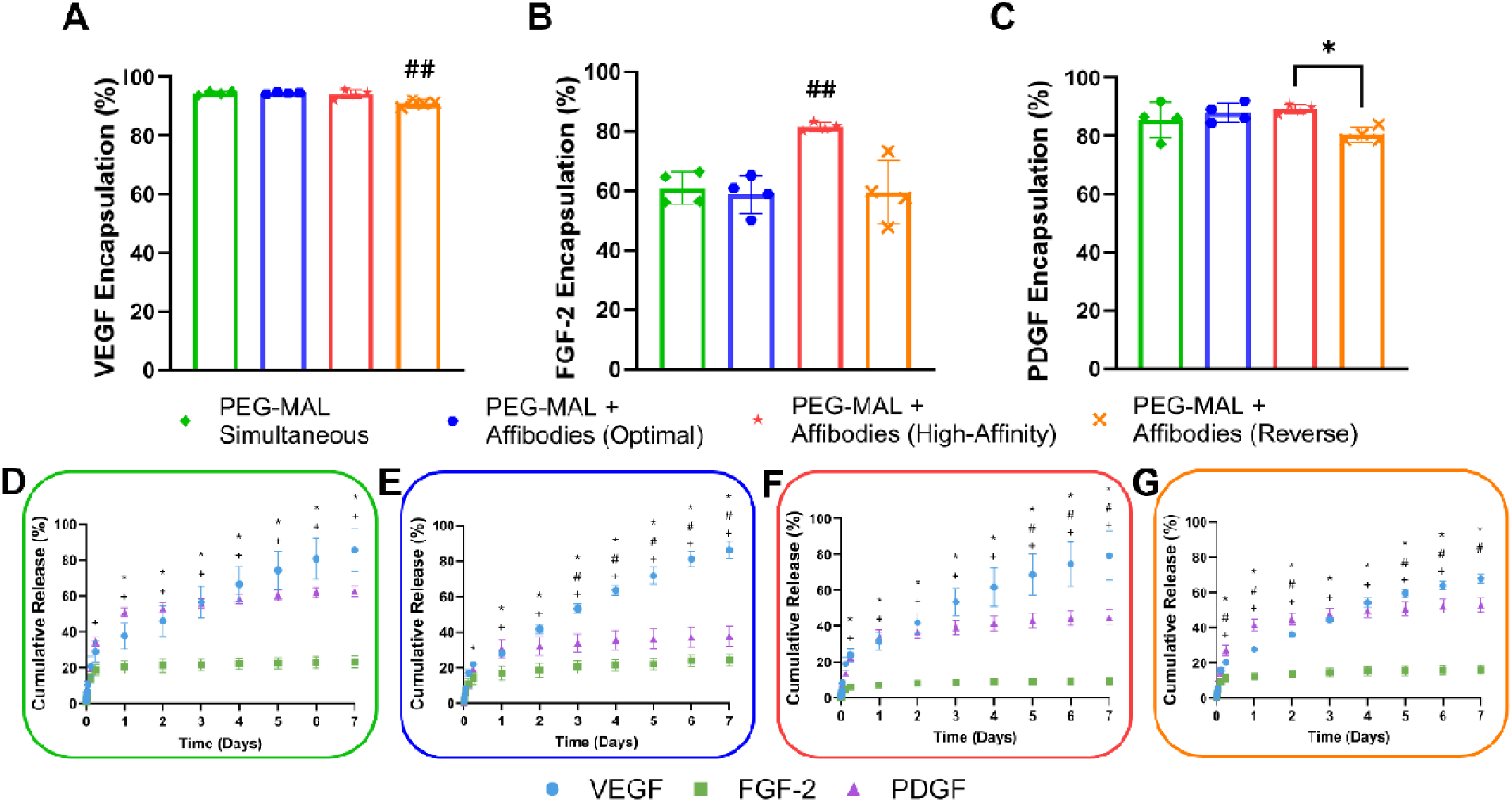
Controlled release of VEGF, FGF-2, and PDGF from multiple affibody-conjugated hydrogels. Encapsulation efficiency of A) VEGF, B) FGF-2, and C) PDGF loaded into PEG-MAL hydrogels containing no affibodies or multiple protein-specific affibodies. Statistical significance determined by one-way ANOVA and Tukey’s post hoc test. (n=4, * p < 0.05, ## significantly different from all other groups) Release of VEGF, FGF-2, and PDGF from D) PEG-MAL Simultaneous, E) PEG-MAL + Affibodies (Optimal), F) PEG-MAL + Affibodies (High Affinity), and G) PEG-MAL + Affibodies (Reverse) hydrogels. Statistical significance determined by two-way ANOVA and Tukey’s post hoc test. (n=4, * p < 0.05, * VEGF vs. FGF-2, # VEGF vs. PDGF, + FGF-2 vs. PDGF)

We next evaluated cumulative release of VEGF, FGF-2, and PDGF from each hydrogel composition over 7 days and graphed these release profiles two different ways to compare release of individual proteins between hydrogel compositions (**Fig. S2**) and to compare release of all target proteins within each hydrogel composition (**Fig. 3D-G**). As expected, the PEG-MAL + Affibodies (Reverse) hydrogels containing the VEGF High Affibody released less VEGF between Days 3 and 7 than the PEG-MAL Simultaneous and PEG-MAL + Affibodies (Optimal) hydrogels, which did not contain VEGF-specific affibodies **(Fig. S2A)**. Interestingly, a lower cumulative release of VEGF was not observed for the PEG-MAL + Affibodies (High-Affinity) hydrogels, which also contained the VEGF High Affibody, contradicting the expected outcome of the VEGF High Affibody reducing VEGF release from the hydrogel. This result may be caused by an increase in non-specific, off-target interactions between the different growth factors and other affibodies in these hydrogels that may interfere with the intended affinity interactions in this complex environment and confined space.

Significant differences were observed for FGF-2 release between all groups starting at Day 1 and persisting until Day 7 except between the PEG-MAL Simultaneous and PEG-MAL + Affibodies (Optimal) hydrogels (**Fig. S2B**). The PEG-MAL + Affibodies (High-Affinity) hydrogels displayed the lowest cumulative FGF-2 release, likely due to the incorporation of the FGF-2 High Affibody resulting in greater FGF-2 retention. The PEG-MAL + Affibodies (Reverse) hydrogels released less FGF-2 than the PEG-MAL + Affibodies (Optimal) hydrogels despite both hydrogels containing the FGF-2 Medium Affibody and displaying no differences in initial FGF-2 encapsulation (**Fig. S2B**). This result may be described by the observed small degree of non-specific binding between the VEGF High Affibody and FGF-2 (**Fig. 1E**), resulting in greater retention of FGF-2 within the PEG-MAL + Affibodies (Reverse) hydrogels.

Significant differences were also observed for PDGF release between all groups between Days 4 and 7 **(Fig. S2C)**. Hydrogels without affibodies released the most PDGF due to the lack of affinity interactions to slow protein release. Conversely, both the PEG-MAL + Affibodies (High-Affinity) and PEG-MAL + Affibodies (Optimal) hydrogels released the least PDGF over 7 days, consistent with the expected impact of incorporating PDGF High Affibodies within these hydrogels. The PEG-MAL + Affibodies (Reverse) hydrogels released less PDGF than the PEG-MAL Simultaneous hydrogels despite both hydrogel compositions lacking PDGF-specific affibodies. This result was likely due to the presence of the FGF-2 Medium Affibody in the PEG-MAL + Affibodies (Reverse) hydrogels, which exhibited some off-target binding to PDGF on BLI (**Fig. 1H**).

We next sought to determine how variations in multiple affibody-conjugated hydrogel compositions impacted the relative release of VEGF, FGF-2, and PDGF. The PEG-MAL Simultaneous hydrogel released the most VEGF, FGF-2, and PDGF over 7 days **(Fig. 3D)**. The PEG-MAL + Affibodies (Optimal) hydrogel released less VEGF, FGF-2, and PDGF than the PEG-MAL Simultaneous hydrogels, with equal amounts of VEGF and PDGF release until Day 2 **(Fig. 3E)**. The PEG-MAL + Affibodies (High-Affinity) hydrogel demonstrated reduced FGF-2 and VEGF release across all timepoints **(Fig. 3F)**. The PEG-MAL + Affibodies (Reverse) composition released more PDGF than VEGF at earlier timepoints, consistent with the expected impact of not containing PDGF-specific affibodies **(Fig. 3G)**. Taken together, these results demonstrate the efficacy of multiple affibody-conjugated hydrogels in temporally regulating the relative abundance of released VEGF, FGF-2, and PDGF.

### Single growth factor delivery does not impact MVF network length and branching

To explore how the timing of VEGF, FGF-2, and PDGF presentation impacts angiogenesis, we first investigated the effect of these three growth factors on HUVEC vascular network formation. However, we observed a limited impact of separate or combined VEGF, FGF-2, and PDGF treatment on HUVEC network length and branching (**Fig. S3A-B**). This result may be described by the limitation of endothelial cell monocultures in exploring the multiple stages of angiogenesis, which involves the coordinated actions of multiple cell types, including pericytes, macrophages, and fibroblasts, which each exhibit unique responses to VEGF, FGF-2, and PDGF treatment.^10,12,16^ To address this limitation, we adopted an *in vitro* multicellular MVF model containing mature vascular fragments and a variety of support cells to probe how temporal variations in the phased presentation of VEGF, FGF-2, and PDGF impact vascular network formation. Rat MVFs were seeded in collagen type I hydrogels and incubated for 3 days prior to treatment with 50 nM of VEGF, FGF-2, or PDGF on Day 3 (**Fig. S4**), 50 nM of all three growth factors on Day 3, or 50 nM of one growth factor per day on Days 3, 4, and 5 **(Fig. 4A)**. MVFs were cultured for 7 days, followed by fixation, lectin staining to visualize vascular structures, and imaging by confocal microscopy. Total vascular network length and branching were quantified and compared to untreated MVFs from each primary tissue harvest to determine the impact of soluble protein treatments on vascular network formation **(Fig. 4B)**. For single growth factor treatments, RT-qPCR was also performed on Day 7 to evaluate endpoint gene expression.

**Figure 4.**
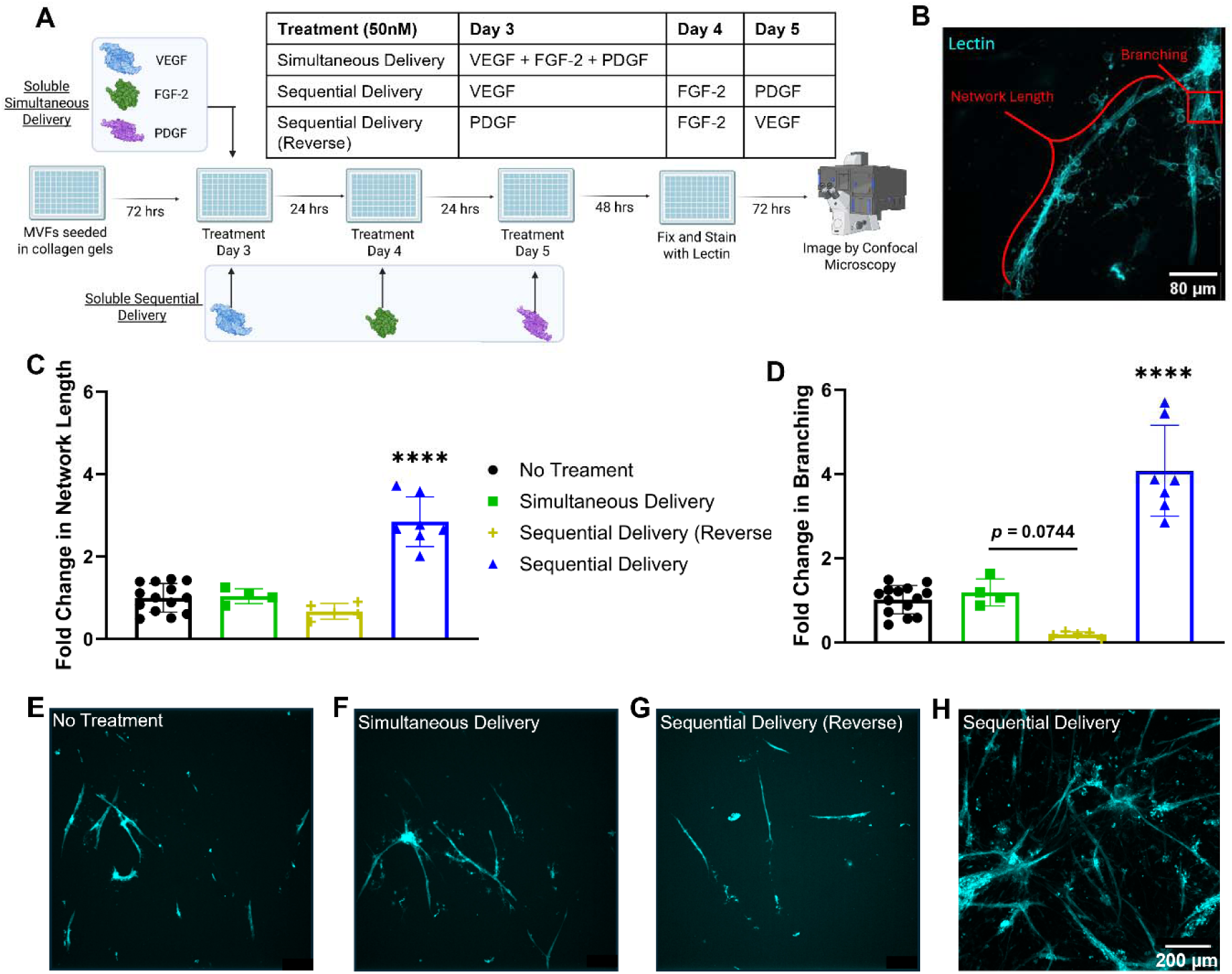
Effect of simultaneous or sequential delivery of soluble VEGF, FGF-2, and PDGF on rat-derived MVF network formation. A) Schematic of simultaneous and sequential delivery of soluble VEGF, FGF-2, and PDGF on days 3, 4, and 5 to MVFs seeded in collagen type I hydrogels. B) Representative lectin-stained maximum intensity z-projection confocal microscopy image of untreated MVFs with branch points and network length highlighted in red. Fold change in Day 7 C) vascular network length and D) branching of treated MVFs normalized to untreated MVFs quantified from confocal microscopy images. Statistical significance was determined by one-way ANOVA and Tukey’s post-hoc test. (n=4-14, ****p < 0.0001 compared to all other groups) Representative lectin-stained maximum intensity z-projection confocal microscopy images of MVFs with E) No Treatment, F) Simultaneous Delivery, G) Sequential Delivery (Reverse), and H) Sequential Delivery. Scale bar = 200 μm.

MVFs treated with single growth factors showed no significant increase in network length or branching compared to untreated MVFs, indicating no impact on vascular network formation (**Fig. S4A-B**). However, significant changes in angiogenic gene expression were observed with each treatment (**Fig. S4C-I**), suggesting that the growth factors each stimulated different cell behaviors. As expected, VEGF treatment increased ANGPT-2 and decreased TIMP-1 expression, suggesting active destabilization of vascular networks. FGF-2 treatment increased ANGPT-2 and MMP-3 expression, suggesting active vessel destabilization and extracellular matrix remodeling. Interestingly, PDGF increased expression of ANGPT-1, ANGPT-2, and MMP-3 and decreased expression of TIMP-1, suggesting concurrent, competing upregulation of vascular network stabilization and extracellular matrix remodeling. Although this result was unexpected due to the canonical role of PDGF in recruiting pericytes to stabilize vascular networks, this outcome may be described by changes in the ratios of endothelial cells, pericytes, fibroblasts, and adipocytes found within the MVF culture, as these cell types all respond to PDGF to different extents.^17,53^ No significant changes were observed in PDGFRΒ, VEGF-A, or VEGFR-2 expression. Collectively, these results suggest that, although single growth factor treatment of MVFs over 4 days was sufficient to change angiogenic gene expression, these changes did not alter the formation of vascular networks.

### Sequential delivery of VEGF, FGF-2, and PDGF increases MVF network length and branching

Sequential delivery of soluble VEGF, followed by FGF-2, and then PDGF in the optimal order expected to typically occur during angiogenesis increased microvascular network length and branching compared to no treatment, simultaneous delivery of all three proteins, and sequential delivery of proteins in the reverse order (**Fig. 4C-D**). A modest decrease in vascular branching (p = 0.0744) was also observed with Sequential Delivery (Reverse) compared to Simultaneous Delivery, indicating that PDGF delivery prior to VEGF and FGF-2 delivery may recruit pericytes that stabilize endothelial cell-cell junctions and inhibit the vascular fenestration required for sprouting.^8,53^ This finding is further supported by soluble PDGF treatment upregulating ANGPT-1 expression, which is associated with stabilization of endothelial cell junctions.^54^ Taken together, these data suggest that phased growth factor secretion may be necessary to coordinate the stages of angiogenesis required for vascular outgrowth and that simultaneous presentation of all three growth factors conversely diminishes vascular network formation through concurrent activation of opposing cellular functions.^8,21,55,56^

### Phased VEGF, FGF-2 and PDGF co-delivery using multiple protein-specific affibodies increases MVF network length and branching

Motivated by the effects of sequential soluble protein delivery on MVF network formation, we formulated affibody-conjugated hydrogels to achieve similar phased release of VEGF, FGF-2, and PDGF to MVFs **(Fig. 5A)** using different combinations of affibodies specific for each growth factor (i.e., PEG-MAL + Affibodies (Optimal), PEG-MAL + Affibodies (High-Affinity), and PEG-MAL + Affibodies (Reverse)) **(Fig. 5B**). Hydrogels were loaded with 17.5 pmol of each growth factor and transferred in transwells to 24-well plates containing MVFs seeded in type I collagen hydrogels.

**Figure 5.**
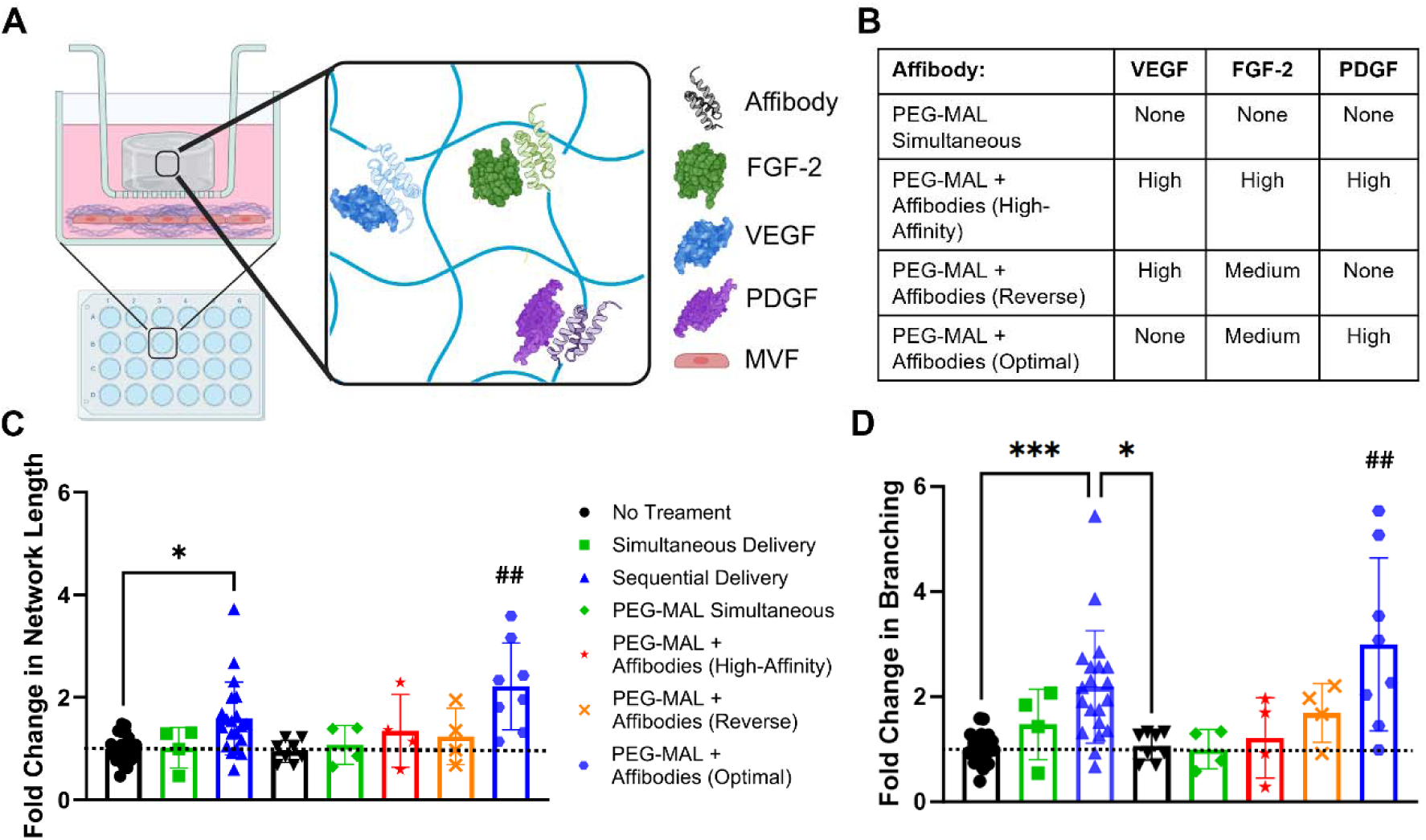
Effect of phased delivery of VEGF, FGF-2, and PDGF from affibody-conjugated PEG-mal hydrogels on rat-derived MVF network formation. A) Schematic of transwell setup with affibody-conjugated PEG-mal hydrogels and type I collagen hydrogels containing MVFs. Affibody-conjugated hydrogels for controlled VEGF, FGF-2, and PDGF delivery are placed in the top compartment, while collagen hydrogels containing MVFs are placed in the bottom compartment. B) Table of protein-specific affibodies used in each hydrogel formulation. Fold change in Day 7 C) vascular network length and D) branching of treated MVFs normalized to untreated MVFs quantified from confocal microscopy images. MVFs were treated with soluble growth factors or growth factors delivered from affibody-conjugated hydrogels in transwell inserts. Statistical significance was determined by one-way ANOVA and Tukey’s post-hoc test. (n=4-19, *p < 0.05, **p < 0.01, ***p < 0.001, ## different from every other group)

Vascular network length and branching for each treatment group was normalized to untreated MVFs. Network length and branching of MVFs treated with soluble VEGF, FGF-2 and PDGF simultaneously or sequentially were consistent between the 96-well and 24-well plate formats. Treatment with the PEG-MAL hydrogel delivery vehicle without growth factors (PEG-MAL), with growth factors without affibodies (PEG-MAL Simultaneous), with growth factors and high affinity affibodies (PEG-MAL + Affibodies (High-Affinity)), and with growth factors and affibodies that provided sequential protein release in the reverse order (PEG-MAL + Affibodies (Reverse)) all had no effect on MVF network length and branching compared to untreated MVFs (**Fig. 5C, D**). In all treatment conditions with growth factors, we expected that modulating protein release with suboptimal timing (i.e., reverse order, fast release, or slow release) would inhibit the formation of robust vascular networks. In particular, for PEG-MAL hydrogels providing reverse protein release, we hypothesized that initial release of PDGF followed by FGF-2 and then VEGF resulted in greater pericyte adherence to the lumen wall and subsequent diminished vascular sprouting needed for vascular network formation. Conversely, treatment with PEG-MAL + Affibodies (Optimal) significantly increased both vascular network length and branching compared to all other treatment groups and untreated MVFs. This result suggests that this multiple affibody-conjugated hydrogel composition successfully phased the release of VEGF, FGF-2, and PDGF to exceed the enhancement of vascular network formation provided by soluble, sequential treatment with these three growth factors in the optimal order.

### Phased VEGF, FGF-2 and PDGF co-delivery using multiple protein-specific affibodies alter the gene expression of multicellular MVF networks

To further understand the gene expression changes responsible for the formation of MVF vascular networks in response to growth factor delivery, we performed RT-qPCR on MVFs treated with soluble and hydrogel-mediated delivery of VEGF, FGF-2, and PDGF. Similar to the gene expression analysis of our single growth factor treatment groups, we evaluated the expression of MMP-3, TIMP-1, ANGPT-1, ANGPT-2, VEFG-A, VEGFR-2, and PDGFRB at Day 7 (**Fig. 6A-G**). We also performed ELISAs for VEGF, FGF-2, and PDGF on MVF-conditioned media from Day 7 to relate the microenvironmental concentrations of VEGF, FGF-2, and PDGF to these gene expression profiles (**Fig. S5A-C**). Overall, soluble delivery of growth factors between Days 3 and 5 resulted in higher growth factor concentrations in the conditioned media at Day 7 compared to hydrogel-mediated release of growth factors starting at Day 3.

**Figure 6.**
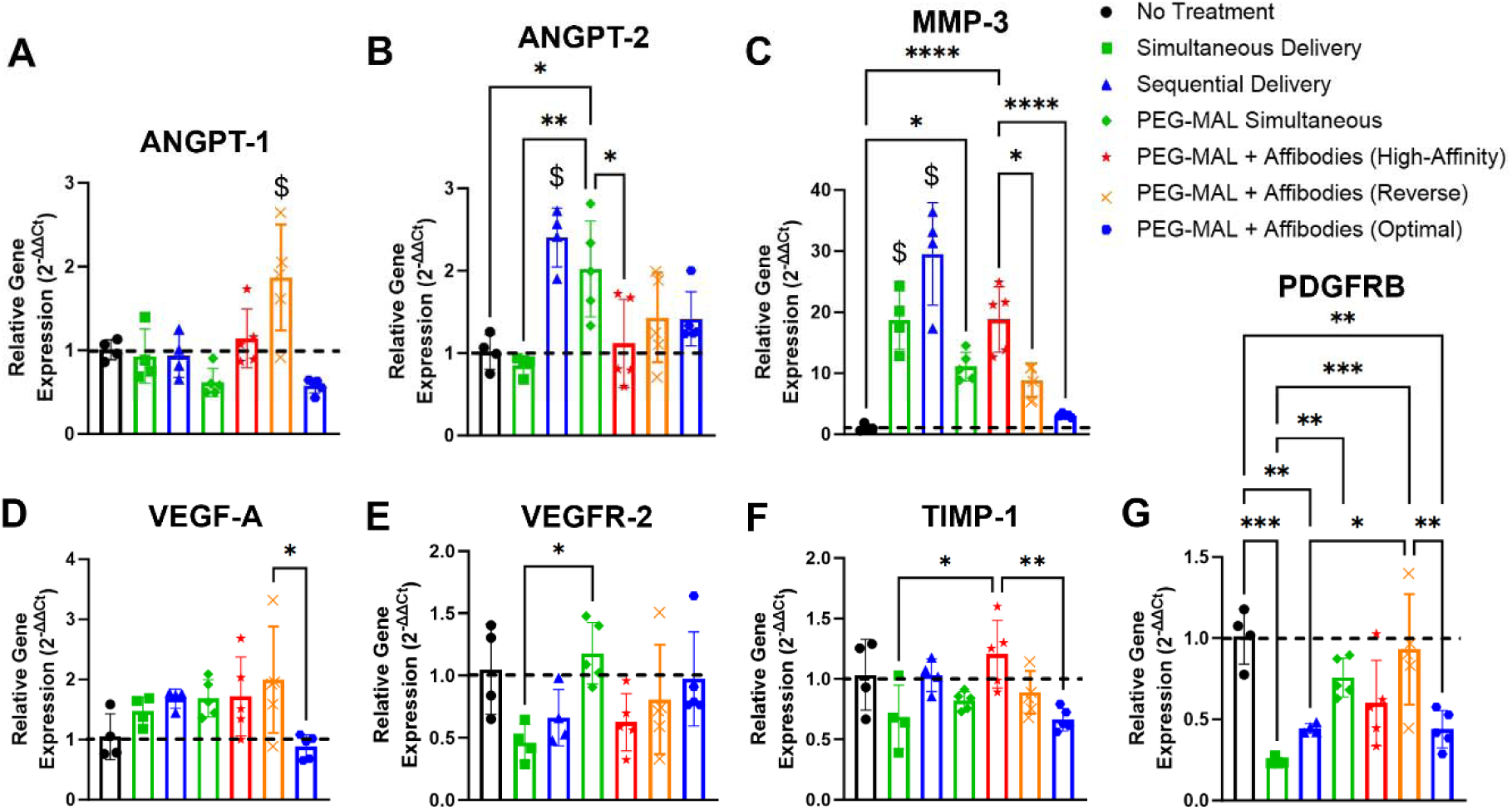
Gene expression of MVFs on Day 7 after simultaneous or phased delivery of VEGF, FGF-2, and PDGF. Relative gene expression of A) ANGPT-1, B) ANGPT-2, C) MMP-3, D) VEGF-A, E) VEGFR-2, F) TIMP-1, and G) PDGFRB were measured by RT-qPCR using TaqMan probes. Data were normalized to ACTB expression using the 2^-ΔΔCt^ method and plotted as fold change compared to untreated MVFs on Day 7. Statistical significance was determined by one-way ANOVA and Tukey’s post-hoc test. (n=4, *p < 0.05, **p < 0.01, ***p < 0.001, ****p < 0.0001, $ different from every other group)

Among soluble growth factor treatments, both Simultaneous Delivery and Sequential Delivery displayed increased MMP-3 and decreased PDGFRB expression at Day 7, suggesting ECM remodeling and a potential decrease in the number of pericytes compared to other cell types within culture (**Fig. 6C,G**). Sequential Delivery also stimulated higher ANGPT-2 expression than all other groups, suggesting the occurrence of increased vascular fenestration and endothelial layer destabilization, key angiogenic processes that align with the vascular network expansion observed (**Fig. 6B**). Interestingly, despite the difference in ANGPT-2 expression and MVF network length and branching, both soluble treatment groups contained similar amounts of each growth factor in the conditioned media at Day 7. Thus, these differences may be explained by differences in the duration of growth factor exposure, as the Simultaneous Delivery treatment condition exposed the MVFs to all three proteins from Day 3 to 7, while the Sequential Delivery treatment condition only exposed the MVFs to VEGF for the same amount of time and to FGF-2 and PDGF for less time.

We next wanted to explore how simultaneous delivery of VEGF, FGF-2, and PDGF from a hydrogel containing no affibodies (PEG-MAL Simultaneous) affected gene expression and growth factor concentrations in MVF cultures after 7 days. In contrast to treatment with soluble growth factors, simultaneous delivery of VEGF, FGF-2, and PDGF from a hydrogel stimulated increased expression of ANGPT-2, VEGFR-2, and PDGFRB compared to soluble Simultaneous Delivery and decreased expression of MMP-3 compared to Simultaneous Delivery (**Fig. 6B,E,G**). Moreover, the ELISA data revealed that the concentrations of VEGF, FGF-2, and PDGF in the conditioned media from PEG-MAL Simultaneous were approximately 4-5-fold lower than the concentrations of these growth factors in the Simultaneous Delivery treatment condition on Day 7 but higher than growth factor concentrations in conditioned media from the affibody-containing hydrogels. These differences suggested that sustained, diffusion-mediated release of the growth factors from a PEG-MAL hydrogel sufficiently altered protein concentrations in the MVF cultures to trigger distinct changes in gene expression indicative of increased vessel destabilization.

Treatment with the PEG-MAL + Affibodies (High-Affinity) hydrogels increased MMP-3 expression compared to no treatment and increased TIMP-1 expression compared to Simultaneous Delivery and treatment with the PEG-MAL + Affibodies (Optimal) hydrogel (**Fig. 6C,F**). Since TIMP-1 inhibits matrix metalloproteases like MMP-3, this combination of increased MMP-3 and TIMP-1 expression may indicate a balance between ECM growth and remodeling by Day 7. Similarly, the PEG-MAL + Affibodies (Reverse) hydrogel was the only condition to increase expression of ANGPT-1 and VEGF-A, which are vascular stabilization and destabilization cues, respectively (**Fig. 6A,D**). These two sets of competing cues may explain why poor vascular network formation was observed with both the PEG-MAL + Affibodies (High Affinity) and PEG-MAL + Affibodies (Reverse) hydrogels (**Fig. 5C-D**). In fact, treatment with both hydrogel compositions resulted in an absence of VEGF in the conditioned media at Day 7. The incorporation of the VEGF High Affibody likely increased VEGF retention within the hydrogel, which may further explain the lack of vascular network formation.

Interestingly, despite the fact that the PEG-MAL + Affibodies (Optimal) hydrogel stimulated significantly higher MVF network length and branching, few differences in gene expression were observed between the PEG-MAL + Affibodies (Optimal) hydrogel and other treatment groups. The most notable differences in gene expression were observed between the PEG-MAL + Affibodies (Optimal) hydrogel and the other affibody-conjugated hydrogels, which suggest that changing the release profiles of the growth factors significantly altered the cellular recruitment, patterning, and maturation required for vessel formation and stabilization. These differentially expressed genes included decreases in ANGPT-1, PDGFRB, MMP-3, VEGF-A, and TIMP-1 expression. Furthermore, we observed a decrease in PDGFRB expression and no significant differences in gene expression of stabilization markers TIMP-1 and ANGPT-1 and destabilization markers MMP-3, ANGPT-2, and VEGF-A between treatment with PEG-MAL + Affibodies (Optimal) and untreated MVFs, suggesting the presence of stable vascular networks on Day 7. In contrast, the significant changes in gene expression in the other treatment groups compared to untreated MVFs may indicate vascular networks that are actively remodeling. Collectively, these gene expression results suggest that vascular networks from the different treatment groups may be at different stages of growth and maturation, explaining the differences in network length and branching that we observed at Day 7, and suggesting that our hydrogel delivery vehicles are differentially affecting specific angiogenic genes within the MVF model.

### Phased VEGF, FGF-2, and PDGF co-delivery from affibody-conjugated hydrogels does not affect perivascular coverage

PDGF secretion has been implicated in stimulating pericyte recruitment to the vessel lumen in later stages of angiogenesis, resulting in stabilization of neovessel sprouts.^16,53,57^ Thus, we next explored the effect of soluble and affibody-mediated PDGF delivery on perivascular coverage of our microvascular networks. After 7 days of culture, we stained MVFs with lectin for visualizing vascular structures and α-smooth muscle actin (α-SMA) for pericytes and quantified co-localization of lectin and α-SMA staining to determine the amount of perivascular coverage in each treatment group, described by the Mander’s overlap coefficient.

Representative images for each treatment all showed some degree of co-localization (pink) between lectin staining (red) and α-SMA staining (blue) (**Fig. 7A-G**). Interestingly, no differences in the Mander’s overlap coefficients were observed between groups despite differences in overall MVF network length and branching (**Fig. 7H**). These results suggest that growth factor delivery either solubly or via affibody-conjugated hydrogels did not dysregulate pericyte function and that treatment with the PEG-MAL + Affibodies (Optimal) hydrogel expanded the pericyte population to sufficiently cover the newly expanded vascular networks. Perivascular coverage coupled with the unchanged vessel stabilization/destabilization gene expression for the PEG-MAL + Affibodies (Optimal) hydrogel provide further support for the theory that the vascular networks observed at Day 7 were stable.

**Figure 7.**
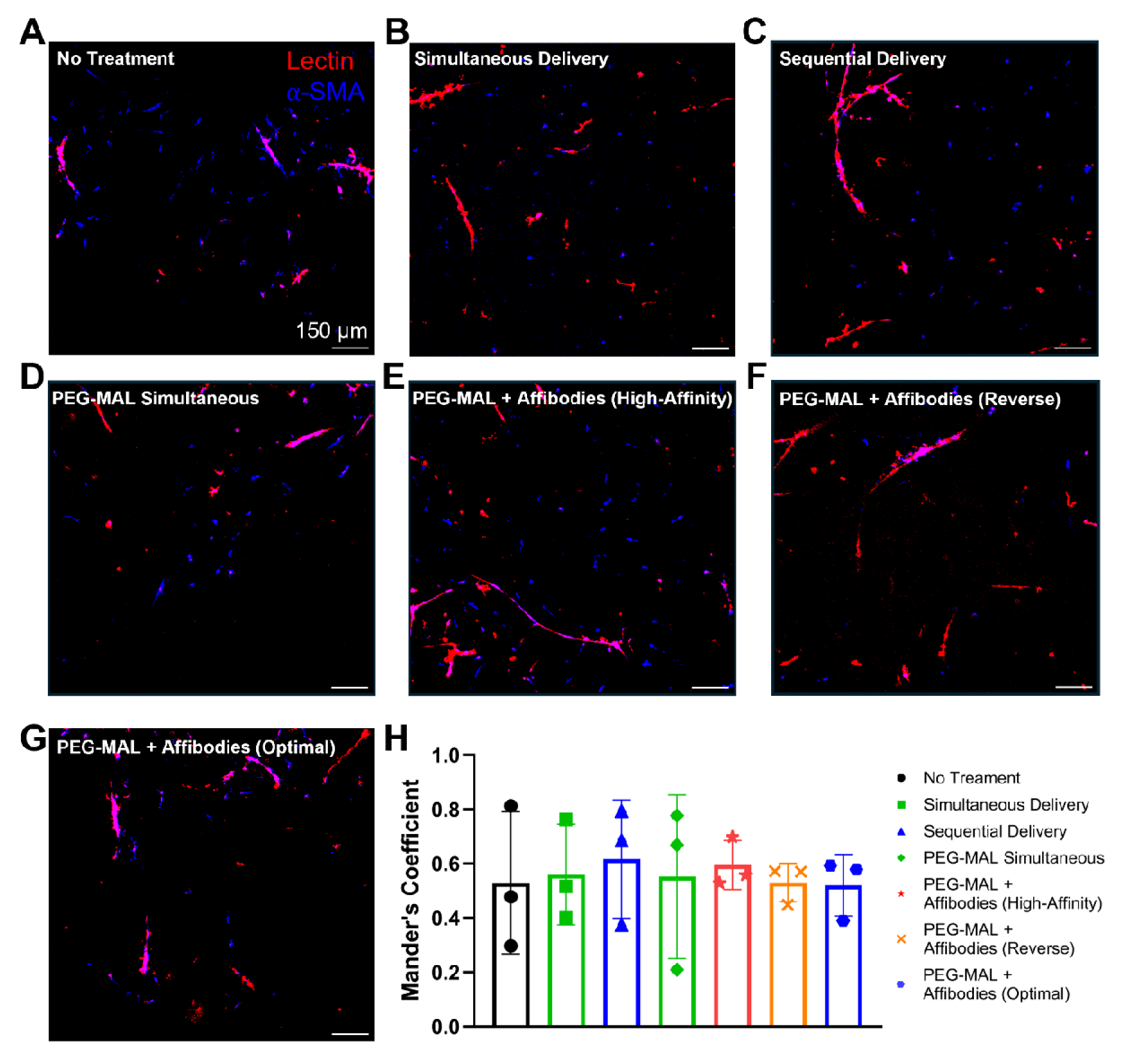
Impact of phased delivery of VEGF, FGF-2, and PDGF on perivascular coverage at Day 7. Single slice representative confocal microscopy images of lectin (red) and α-SMA (blue) stained MVFs with overlapping signal depicted in pink. MVFs received A) no treatment, B) simultaneous delivery of soluble VEGF, FGF-2, and PDGF, C) sequential, soluble delivery of VEGF followed by FGF-2 and PDGF, D) simultaneous delivery using PEG-MAL hydrogels, or E-G) varying phased delivery of VEGF, FGF-2, and PDGF using different affibody-conjugated PEG-MAL hydrogels. Scale bar = 150 µm. H) Mander’s overlap coefficient for lectin and α-SMA staining colocalization.

## Discussion

Angiogenesis requires the temporally regulated presentation of key signaling proteins to stimulate distinct cell behaviors that contribute to higher order cell patterning that is inherent to mature vasculature. Dysregulation in protein secretion is notoriously difficult to study within complex *in vivo* environments. *In vitro* platforms present opportunities to study how variations in secreted morphogen gradients impact cell signaling. Protein delivery vehicles capable of predictably altering the environmental concentrations of multiple proteins allow us to explore the role of temporal protein presentation in mediating the stages of angiogenesis. Affinity-based delivery systems are well suited for this approach, as affibodies provide an easily tunable, protein-specific, and material-independent mechanism for regulating protein delivery. The highly modular and customizable nature of using different affibodies to control the release rates of different proteins provides the advantage of offering better tunability and precision than other protein release approaches that rely upon material degradation or non-specific polymer-protein interactions to control protein release.^5,22,58^ Previous work has demonstrated that phased VEGF, FGF-2, and PDGF delivery triggered by scaffold degradation stimulates the formation of robust vascular networks from HUVECs and chorioallantoic membranes *in vitro*.^5^ Our results corroborate the results of this and other studies,^6,59,60^ demonstrating that similar phased delivery can be achieved using affibody-conjugated hydrogels and that these hydrogel delivery vehicles can direct angiogenesis better than single growth factor delivery or multiple growth factor delivery. Building further upon this work, we also explored how sub-optimal protein release profiles, such as reverse phased delivery (PDGF, FGF-2, then VEGF) and low protein release, impacted MVF networks – an investigation that was only made possible by the modular nature of affibody addition. Thus, our system enables facile investigation into different sequences of protein delivery, allowing us to gain deeper insights into the role of temporally regulated protein presentation on regenerative processes. Furthermore, our results provide compelling evidence for the efficacy of multiple affibody-functionalized drug delivery vehicles in directing cellular responses and multicellular organization.

In the development of our phased angiogenic growth factor delivery platform, we not only show that phased protein delivery in the optimal sequence significantly increases MVF network length and branching while maintaining perivascular coverage, but that it also modulates angiogenic gene expression and microenvironmental protein concentrations. Surprisingly, this enhanced MVF network was observed with significantly lower protein concentrations than soluble protein delivery, indicating that affibody-mediated protein release may have advantages for protein bioactivity and bioavailability and further suggesting optimal concentration ranges of these morphogens for stimulating vascular outgrowth.

Furthermore, the differences in gene expression observed between the soluble, sequential delivery of VEGF, FGF-2, and then PDGF compared to hydrogel-mediated delivery of this same sequence of growth factors may be explained by the sustained presentation of all three growth factors simultaneously at the outset of the treatment with the hydrogel matching the native secretion profiles of these growth factors within vascular networks actively undergoing angiogenesis *in vivo*. This conjecture has been corroborated by others who have explored how variations in the phased presentation of VEGF, FGF-2, and/or PDGF impacts vascular network formation. For example, co-delivery of soluble PDGF and FGF-2, but not VEGF and FGF-2 or PDGF and VEGF, stimulated the formation of stable vascular networks, highlighting how simultaneous delivery of growth factors with opposing cellular functions may inhibit angiogenesis.^61^ Soluble FGF-2 treatment followed by PDGF treatment has also been shown to enhance vascular network formation compared to simultaneous FGF-2 and PDGF delivery or treatment with PDGF followed by FGF-2.^59^ This work provides deeper insight into the differential impact of phased FGF-2 and PDGF delivery as temporal regulators of angiogenesis. Consistent with previous literature, our findings highlight how growth factors with complementary or opposing cellular functions require phased presentation to coordinate the stages of angiogenesis. Lastly, although outside of the scope of the current work, interactions of these three angiogenic proteins with each other could be investigated in light of the minimal off-target binding of the affibodies observed herein, the structural similarities of the proteins, and the documented cross-family signaling between PDGF and VEGF receptors.^62^ Such off-target interactions could further explain discrepancies observed in growth factor release and changes in gene expression in the presence of all three growth factors.

## Conclusion

Affinity-based protein-protein interactions provide an attractive approach for regulating the individual release profiles of multiple proteins from a single drug delivery vehicle, allowing us to better explore how variations in protein presentation impact regenerative cell signaling processes. Here, we demonstrate that variable affinity binders specific to the angiogenic growth factors VEGF, FGF-2, and PDGF are capable of modulating protein release to direct vascular network formation in a MVF model. Single affibody-conjugated hydrogels demonstrated affinity-mediated release of their target proteins with cumulative release profiles inversely correlated to the affinities of the conjugated affibodies. In hydrogels containing multiple affibodies, affinity-based protein release was largely consistent with protein release observed with single affibody-conjugated hydrogels with some differences, which were likely due to off-target binding interactions between protein-specific affibodies within the same hydrogel. Temporal variations in the soluble delivery of VEGF, FGF-2, and PDGF dramatically impacted angiogenic gene expression and *in vitro* vascular network formation with the sequential delivery of VEGF, followed by FGF-2 and then PDGF increasing vascular network length and branching. We fabricated affibody-conjugated PEG-MAL hydrogels to mimic this sequence of protein delivery, demonstrating that phased, affibody-mediated delivery of VEGF, FGF-2, and PDGF modulated angiogenic gene expression and increased vascular network length and branching beyond that of all other treatments groups, including the soluble sequential protein delivery, despite low concentrations of all three growth factors in the MVF-conditioned media after 7 days of culture. Our work establishing a delivery platform capable of independently tuning the delivery of multiple angiogenic growth factors provides a proof of concept for the use of affibody-functionalized hydrogels for exploring how temporal changes in protein delivery impacts angiogenesis.

The modularity of this platform, via the inclusion of multiple protein-specific affinity interactions, may permit the exploration of other regenerative processes with therapeutic relevance, contributing to our understanding of the temporal regulation of tissue regeneration as a whole.

## Author Contributions

Conceptualization: J.E.S., C.L.A., S.R.N., R.E.G., M.H.H., Data curation: J.E.S., C.L.A., S.R.N., M.R.F., A.H., Formal analysis: J.E.S., C.L.A., S.R.N., A.H., Funding acquisition: M.H.H., J.E.S., R.E.G.; Investigation: J.E.S., C.L.A., S.R.N., J.R.O., M.R.F., A.H., S.C.O., H.B.H, Methodology: J.E.S, C.L.A, A.H., S.R.N, Project administration: J.E.S., C.L.A., R.E.G., M.H.H., Resources: M.H.H., R.E.G., Supervision: M.H.H., R.E.G., Validation: J.E.S., C.L.A., S.R.N., Roles/Writing - original draft: J.E.S., C.L.A., A.H., M.H.H., and Writing - review & editing: M.H.H., R.E.G., J.E.S., C.L.A., S.R.N., J.R.O., M.R.F., A.H.

## Acknowledgements

We are grateful to members of the Hettiaratchi lab for their thoughtful review of this manuscript and to Josh Kupfer for his assistance with MVF isolation. This work was supported by an Oregon Health & Science University (OHSU) Medical Research Foundation New Investigator Grant, a R21 Trailblazer Award and R35 Maximizing Investigators’ Research Award from the National Institutes of Health (NIH) (R21EB032112, R35GM147507), and the Lary Simpson Professorship to M.H.H. J.E.S. is currently supported by an NIH Ruth L. Kirschstein Predoctoral Individual National Research Service Award (F31HL176164) and was previously supported by an NSF Research Traineeship in Molecular Probes and Sensors for Complex Environments (2022168). S.R.N. was supported by the Wu Tsai Human Performance Alliance. J.R.S. was supported by the University of Oregon Molecular Biology and Biophysics Training Program funded by the NIH (T32GM007759). M.R.F. was partly supported by the University of Oregon Summer Program for Undergraduate Research (SPUR), Knight Campus Undergraduate Scholars (KCUS) Program, and University of Oregon Vice President for Research and Innovation (VPRI) Undergraduate Fellowship. A.H. was partially supported by the Knight Campus Undergraduate Scholars (KCUS) Program. Schematic figures were created using BioRender.com.

