## Supplementary Document for "Phased affinity-controlled delivery of vascular endothelial growth factor, fibroblast growth factor-2, and platelet derived growth factor enhances in vitro angiogenesis"

**Supplemental Information**

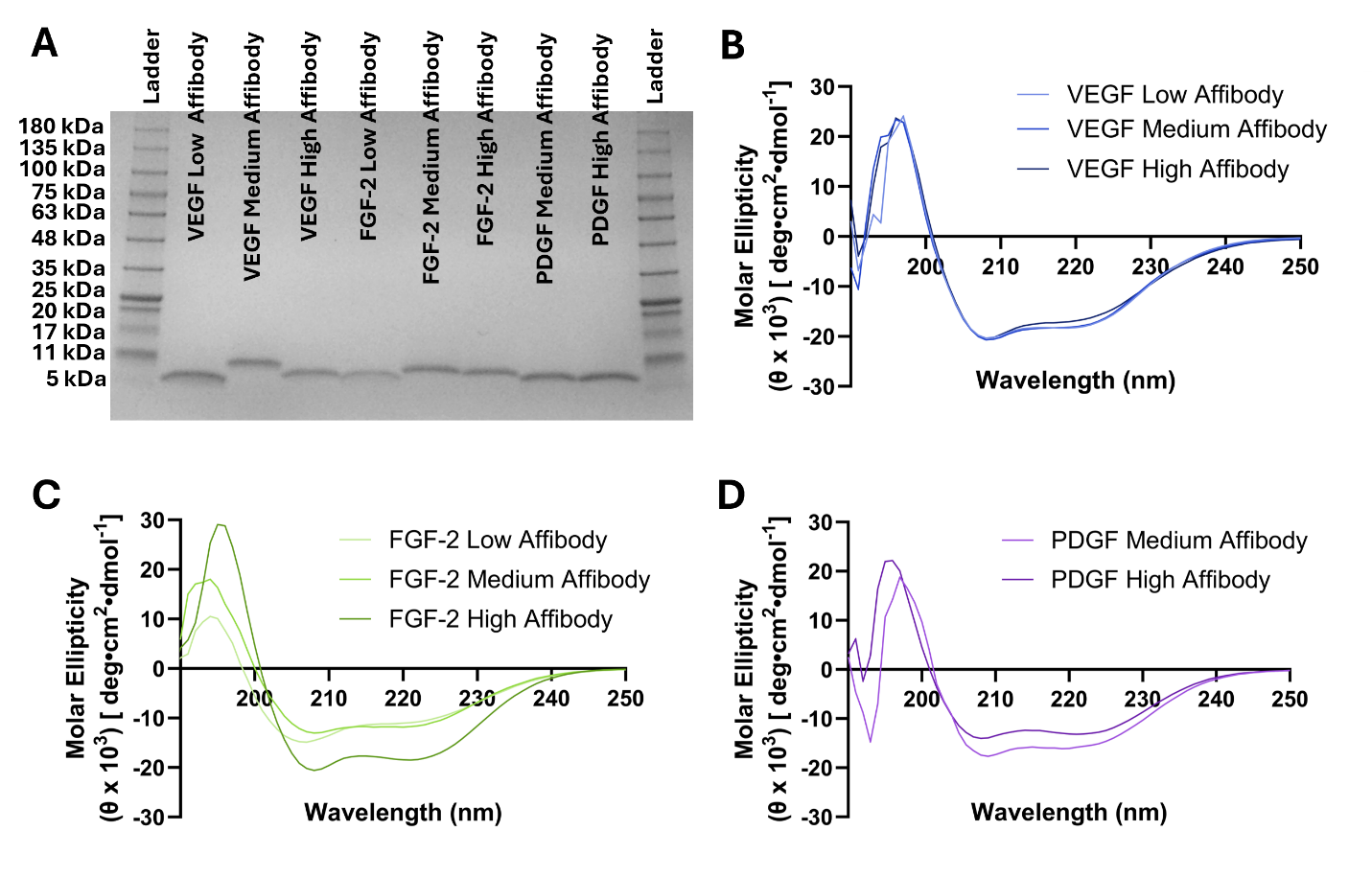

**Figure S1. Biochemical characterization of VEGF-, FGF-2-, and PDGF-specific affibodies.** A) Sodium dodecyl-sulfate polyacrylamide gel electrophoresis (SDS-PAGE) of approximately 2.5 µg of VEGF, FGF-2, and PDGF affibodies compared to a 5-245 kDa reference ladder. Gel was stained using Coomassie Brillant Blue. Circular dichroism spectra of B) VEGF- C) FGF-2- and D) PDGF-specific affibodies measured over a 190-250 nm in 5 mM Tris pH 7.4 at 20 ºC.

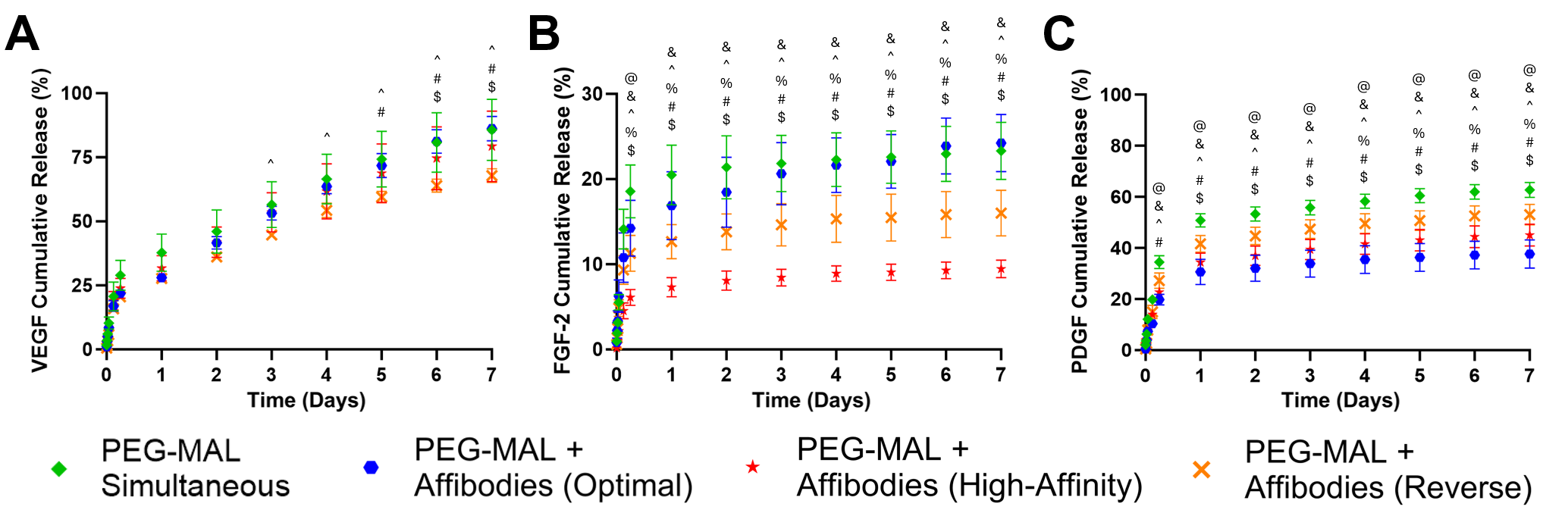

**Figure S2. Controlled release of VEGF, FGF-2, and PDGF from multiple affibody-conjugated hydrogels**. Multiple affibody-conjugated hydrogels for controlled release of A) VEGF, B) FGF-2, and C) PDGF into MVF culture media at 37 °C over 7 days. Statistical significance determined by two-way ANOVA and Tukey’s post hoc test. (n=4, *p < 0.05, @ PEG-MAL Simultaneous vs. PEG-MAL + Affibodies (Optimal), & PEG-MAL Simultaneous vs. PEG-MAL + Affibodies (High-Affinity), ^ PEG-MAL Simultaneous vs. PEG-MAL + Affibodies (Reverse), % PEG-MAL + Affibodies (Optimal) vs. PEG-MAL + Affibodies (High-Affinity), # PEG-MAL + Affibodies (Optimal) vs. PEG-MAL + Affibodies (Reverse), $ PEG-MAL + Affibodies (High-Affinity) vs. PEG-MAL + Affibodies (Reverse))

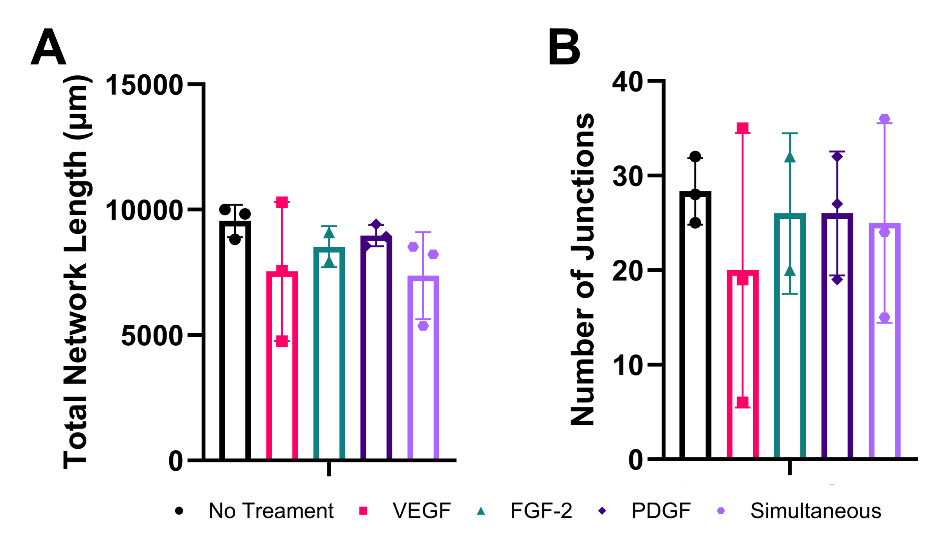

**Figure S3. Impact of delivery of VEGF, FGF-2, and PDGF on HUVEC network formation.** HUVECs were treated with no growth factors, VEGF, FGF-2, PDGF, or all three growth factors simultaneously at 10 ng/mL for 48 hours. ImageJ with the Angiogenesis Analyzer plugin was used to quantify A) total network length and B) number of junctions in an endothelial tube formation assay in response to soluble growth factor delivery. Statistical significance was determined by one-way ANOVA and Tukey’s post-hoc test, and no significant differences were observed. (n=2-3)

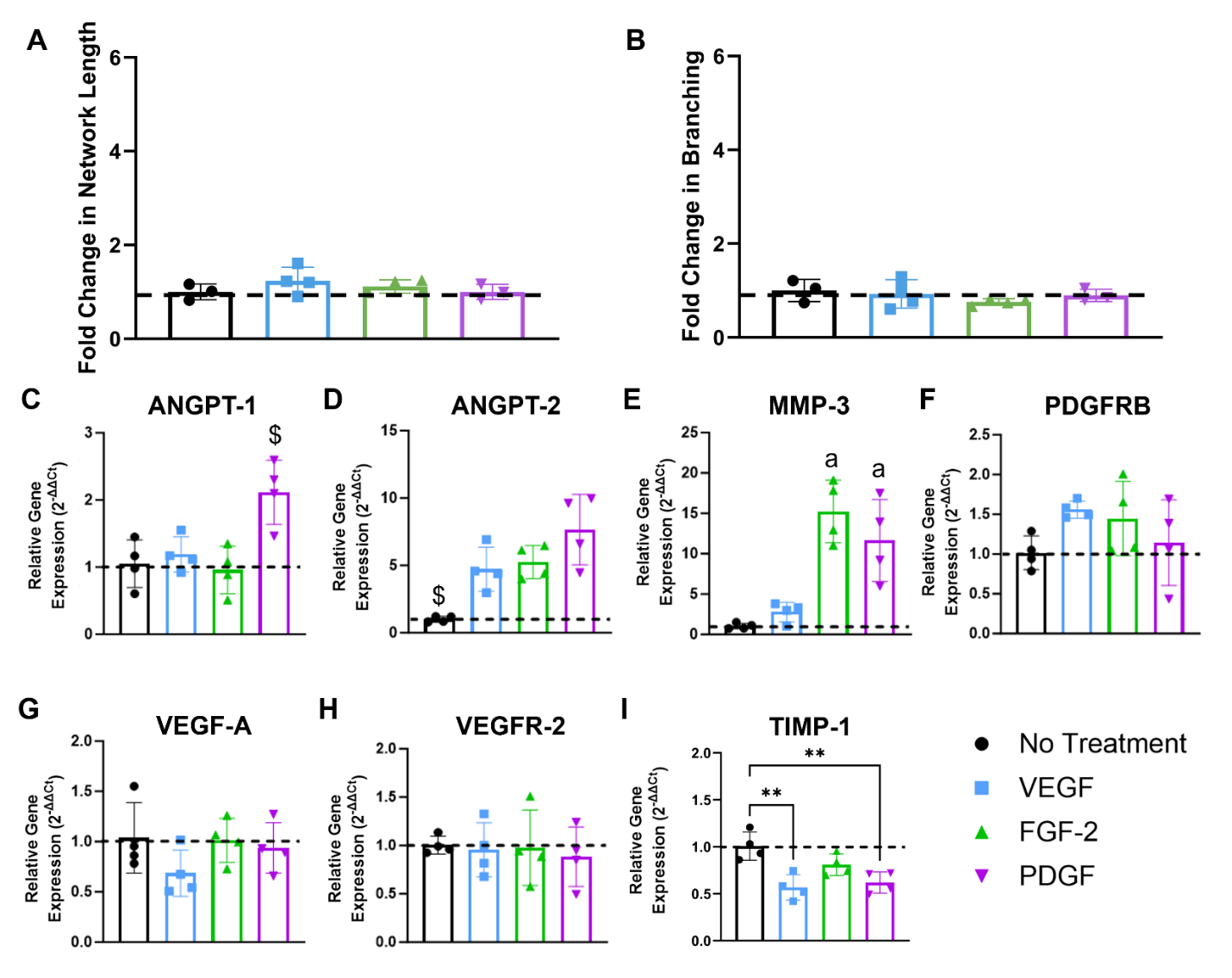

**Figure S4. Impact of VEGF, FGF-2, or PDGF treatment on MVF network length, branching, and angiogenic gene expression.** Fold change in A) vascular network length and B) branching of MVFs treated with either VEGF, FGF-2, or PDGF normalized to untreated MVFs quantified from confocal microscopy images. Statistical significance was determined by one-way ANOVA and Tukey’s post-hoc test. (n = 3-4). Relative gene expression of C) ANGPT-1, D) ANGPT-2, E) MMP-3, F) PDGFRB, G) VEGF-A, H) VEGFR-2, and I) TIMP-1. Data were normalized to ACTB expression using the 2^-∆∆Ct^ method and plotted as fold change compared to untreated MVFs. Statistical significance was determined by one-way ANOVA and Tukey’s post-hoc test. (n=4, **p < 0.01, $ different from every other group, “a” different from every group except each other)

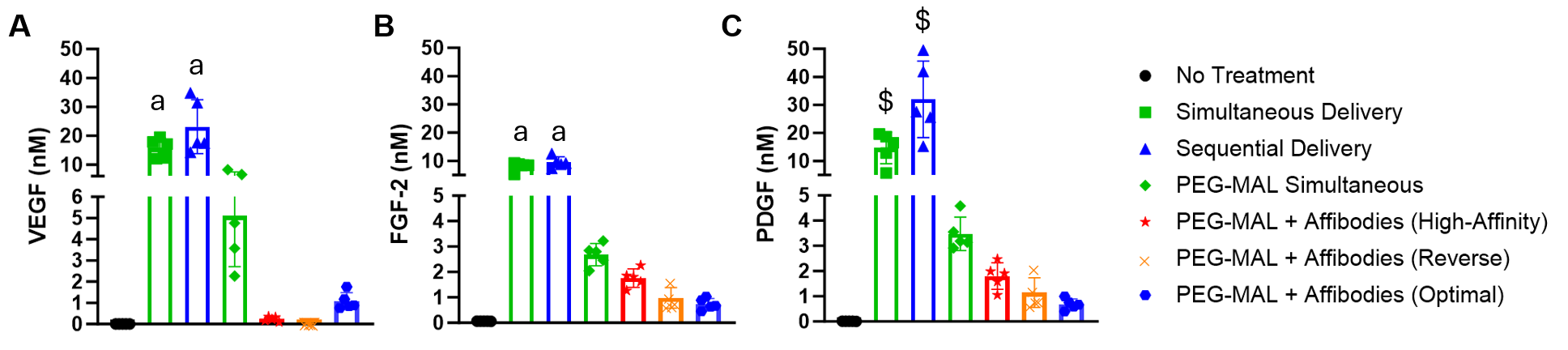

**Figure S5.** **Concentrations of VEGF, FGF-2, and PDGF in Day 7 MVF-conditioned media.** Conditioned media were collected from MVFs at Day 7 prior to collagen hydrogel digestion for RT-qPCR on MVF networks. Concentrations of A) VEGF, B) FGF-2, and C) PDGF following simultaneous or phased delivery of soluble growth factors or growth factors within affibody-conjugated PEG-MAL hydrogels. Statistical significance was determined by one-way ANOVA and Tukey’s post-hoc test. (n=4, “a” different from every group except each other, $ different from every other group)
